# A Hierarchical Bayesian Agent-Based Model for Binary Spatio-Temporal Spread: Theory, PDE Scaling Limit, and an Application to Predator–Prey Cycles

**DOI:** 10.64898/2026.06.03.729943

**Authors:** Xulin Pan

## Abstract

We describe a statistical agent-based model (SABM) for binary spatio-temporal data in which the occupancy of each cell evolves as a Bernoulli mixture of three mechanistically distinct processes: local persistence, anisotropic neighborhood dispersal, and long-distance dispersal. The model is embedded in a hierarchical Bayesian framework with conjugate Beta full-conditionals for the persistence and long-distance parameters and a Dirichlet prior on the directional dispersal kernel. A nonstationary extension links the dispersal kernel to a latent habitat-suitability surface through directional gradients of a Gaussian process. We show that, in the small-step regime, the Lagrangian recurrence for the dispersal kernel scales to a classical two-dimensional advection–diffusion partial differential equation whose drift and dispersion coefficients are the first and second moments of the dispersal probabilities. We provide an MCMC algorithm exploiting the exact full-conditionals and demonstrate parameter recovery and PDE-scaling agreement in a simulated example.

## 1 Introduction

Tracking the spatio-temporal spread of a process—an invasive species, an emerging disease, a wild-fire perimeter—often produces binary data: a cell either is or is not occupied at each sampling time. Mechanistic models of spread (cellular automata, reaction–diffusion equations, agent-based simulators) are expressive but typically lack a likelihood, while purely statistical spatio-temporal models can fit the data but offer little mechanistic interpretation. A *statistical* agent-based model (SABM) bridges these by writing a tractable likelihood directly in terms of interpretable transition probabilities (Hooten and Wikle, 2008; Wikle, 2003; Cressie and Wikle, 2011; Hooten et al., 2020). The approach extends a longer tradition of hierarchical Bayesian dynamical spatio-temporal modeling (Berliner, 1996; Wikle and Hooten, 2010) and complements likelihood-free calibration of mechanistic simulators (Beaumont, 2010; Banks et al., 2017).

We work with a Bernoulli observation model in which the success probability is partitioned into three additive mechanisms: persistence, anisotropic neighborhood dispersal, and long-distance dispersal. The decomposition is exact when the indicator variables identifying each cell’s regime are mutually exclusive, and yields exact conjugate full-conditionals for two of the three parameters. The third—an eight-dimensional direction-resolved dispersal kernel—is modeled with a Dirichlet prior and updated by Metropolis–Hastings. A nonstationary extension lets the dispersal kernel vary across space through gradients of an underlying habitat-suitability Gaussian process, yielding a fully Bayesian hierarchical model that is still amenable to component-wise sampling.

Figure 1 shows the directed acyclic graph that summarizes the hierarchical structure: hyperparameters of the log-Gaussian prior on the kernel concentration and of the Beta priors on *ϕ, ψ* at the top, the latent suitability surface ***α*** and direction-mass log ***a*** in the middle, the cell-specific kernel ***p***_*i*_ as a deterministic function of those latents, and the observed binary occupancy *y*_*i,t*_ at the bottom.

**Figure 1.**
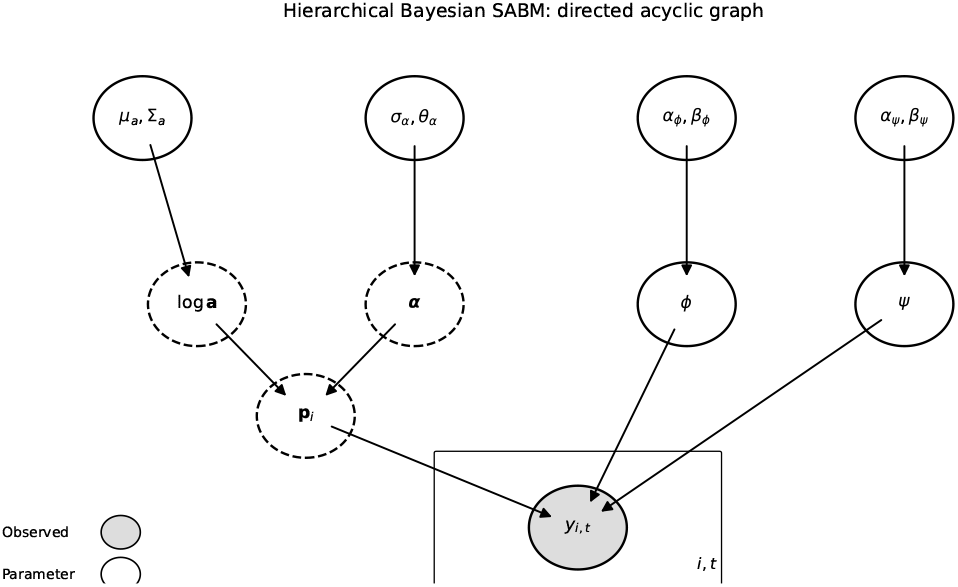
Directed acyclic graph for the hierarchical Bayesian SABM. Shaded node is observed; solid circles are sampled parameters; dashed circles are latent variables. The plate indicates the index (*i, t*) over which *y*_*i,t*_ is repeated.

**Figure 2.**
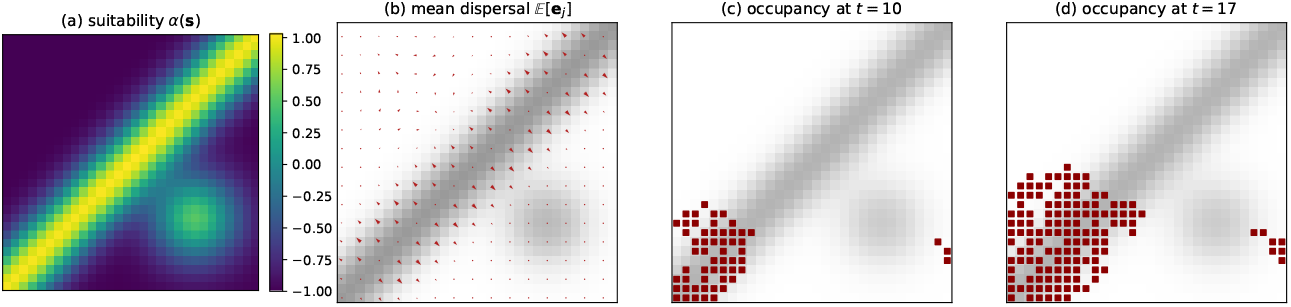
Nonstationary SABM on a 30×30 heterogeneous landscape with a NE-trending suitability ridge and a secondary high-suitability blob. (a) Suitability surface *α*(***s***). (b) Mean dispersal direction ∑_*j*_ *p*_*i,j*_***e***_*j*_ at every cell (arrows), overlaid on *α* in gray. (c,d) Simulated occupancy at *t* = 10 and *t* = 17 from a single seed in the SW corner. The front follows the suitability ridge, then spills into the secondary high-suitability region.

The remainder of the paper is organized as follows. Section 2 states the core data model. Section 3 treats the anisotropic stationary case; Section 4 extends to nonstationary dispersal through habitat suitability. Section 5 gives the full-conditional distributions and the MCMC algorithm. Section 6 derives the advection–diffusion PDE limit. Section 7 demonstrates parameter recovery and PDE agreement in a simulated example.

## 2 Core Data Model

Let *y*_*i,t*_ ∈ {0, 1} denote the occupancy of cell *i* at time *t*, for *i* = 1, …, *N* and *t* = 1, …, *T*. The likelihood is

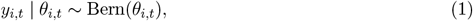

With

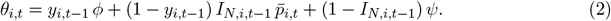

Here

- *ϕ* ∈ (0, 1) is the probability of *persistence* (remaining occupied given currently occupied);
- *ψ* ∈ (0, 1) is the probability of *long-distance dispersal* (becoming occupied when no neighbor is occupied);
- *I*_*N,i,t*−1_ = **1**{∃ *j* ∈ *N*_*i*_ : *y*_*j,t*−1_ = 1} indicates whether at least one neighbor in the neighborhood *N*_*i*_ was occupied at the previous step;
- 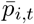 is the probability of *neighborhood-based dispersal*.

The three cases—“occupied,” “vacant with at least one occupied neighbor,” “vacant with no occupied neighbor”—are disjoint and exhaustive, so (2) is a valid probability for every realization of *y*_*i,t*−1_ and 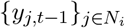.

### Neighborhood dispersal as union of independent arrivals

Let *N*_*i*_ = {*N*_1,*i*_, …, *N*_8,*i*_} be the eight-cell Moore neighborhood of cell *i*, and let 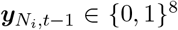 be the vector of neighbor occupancies. Suppose each occupied neighbor in direction *j* contributes an independent “arrival” to cell *i* with probability *p*_*j*_. The probability that *no* neighbor delivers an arrival is 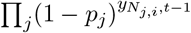, so the probability of at least one arrival is

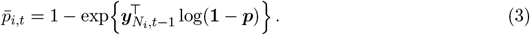

This is the union-of-Bernoullis form: 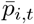 equals *p*_*j*_ when only the direction-*j* neighbor is occupied, and it saturates toward 1 as additional occupied neighbors contribute.

## 3 Anisotropic Stationary Model

In a homogeneous environment the directional probabilities are constant in space. We place a Dirichlet prior on the dispersal kernel,

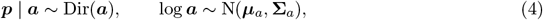

where 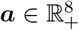 controls the relative weight of each direction. The log-Gaussian prior on ***a*** regularizes the Dirichlet concentrations while allowing strong anisotropy.

## 4 Anisotropic Nonstationary Model

When the environment is heterogeneous, dispersal probabilities should vary spatially. We let an unobserved suitability surface ***α*** ∈ ℝ^*N*^ drive the local kernel:

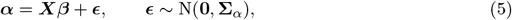

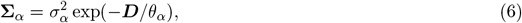

with ***X*** a matrix of spatial covariates, ***β*** unknown coefficients, and ***D*** the matrix of Euclidean distances among cell centroids. The local kernel responds to *gradients* of suitability: agents prefer to move toward more suitable cells. We approximate the gradient at cell *i* in direction *j* by the finite difference

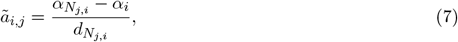

where 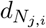 is the Euclidean distance between cell centroids (1 for cardinal directions, 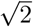 for diagonals). The Dirichlet concentration vector is then

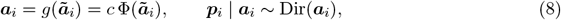

where Φ is the standard-normal CDF (applied componentwise), and *c >* 0 is a scale that controls the overall concentration of the Dirichlet.

The intuition is direct: directions of positive gradient (*ã*_*i,j*_ *>* 0) get Φ(*ã*_*i,j*_) *>* 1*/*2 and hence receive more probability mass; directions of negative gradient are penalized. When the gradient vanishes the distribution is uniform over directions with concentration *c/*2 per arm.

## 5 Inference

### 5.1 Full-conditionals for persistence and long-distance dispersal

Because the three regimes in (2) are disjoint, the likelihood contributions of *ϕ* and *ψ* factor across cells. Let

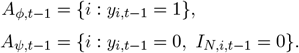

With Beta(*α*_*ϕ*_, *β*_*ϕ*_) and Beta(*α*_*ψ*_, *β*_*ψ*_) priors, the full conditionals are conjugate:

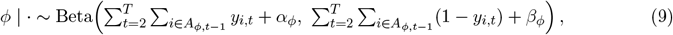

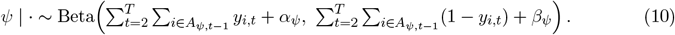

These exact draws hold for both the stationary and the nonstationary submodels.

### 5.2 Updating the dispersal kernel

The neighborhood likelihood does not factor across directions—the union form (3) couples them—so ***p*** (or ***β*** in the nonstationary case) does not enjoy a closed-form full-conditional. We use Gaussian random-walk Metropolis–Hastings updates on log ***a*** (stationary) and ***β*** (nonstationary). With proposal ***a***^*^ = exp(log ***a*** + ***η***), ***η*** ~ N(**0**, *τ* ^2^***I***), the acceptance ratio is

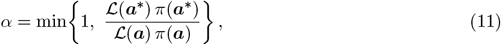

where ℒ is the full ABM likelihood under (1)–(3).

### 5.3 Algorithm

**Algorithm 1 (One MCMC sweep, stationary SABM)**.

1. Draw *ϕ* from (9).
2. Draw *ψ* from (10).
3. Propose log ***a***^*\**^ = log ***a*** + ***η, η*** ~ N(**0**, *τ* ^2^***I***).
4. Compute 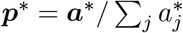 and the proposed log-likelihood *𝓁*^*\**^ = logℒ(*ϕ, ψ*, ***p***^*\**^).
5. Accept with probability min{1, exp(*𝓁*^*\**^ − *𝓁* + log *π*(***a***^*\**^) − log *π*(***a***))}.
6. Return (*ϕ, ψ*, ***a, p***).

For the nonstationary model the third step proposes ***β*** instead, then rebuilds ***α,ã***, ***a***_*i*_, and ***p***_*i*_ via (5)–(8).

## 6 PDE Scaling Limit

The Lagrangian view of the model is that an agent at cell *i* at time *t* moves to cell *N*_*j,i*_ at time *t* + 1 with probability *p*_*j*_. Let 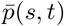 denote the position density of a single such agent. The discrete master equation is

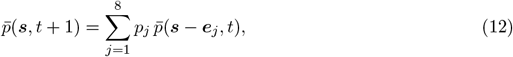

where ***e***_*j*_ ∈ ℝ^2^ is the unit displacement associated with direction *j*. Taylor-expanding 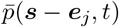 about ***s*** and matching moments gives, in the small-step limit,

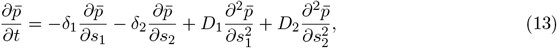

with drift and dispersion coefficients

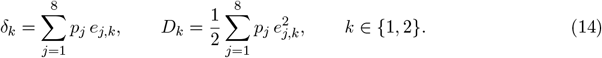

Equation (13) is a classical anisotropic advection–diffusion equation of the kind long used to model biological spread (Skellam, 1951; Okubo and Levin, 2001; Holmes et al., 1994). The mechanistic interpretation is clean: directional bias in ***p*** produces drift, while the spread of the kernel produces dispersion. The same relations (14) link any nonstationary kernel ***p***_*i*_ to spatially varying drift and dispersion fields ***δ***(***s***) and ***D***(***s***), placing the SABM in correspondence with classical reaction– advection–diffusion models of spread. Figure 3 visualizes the two moments for the NE-biased kernel used in the simulation study: the directional probabilities (left) give an overall drift ***δ*** ≈ (0.46, 0.28), and the dispersion coefficients ***D*** ≈ (0.29, 0.21) produce diffusion ellipses (right) whose centres advance along the drift line and whose widths grow as 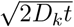.

**Figure 3.**
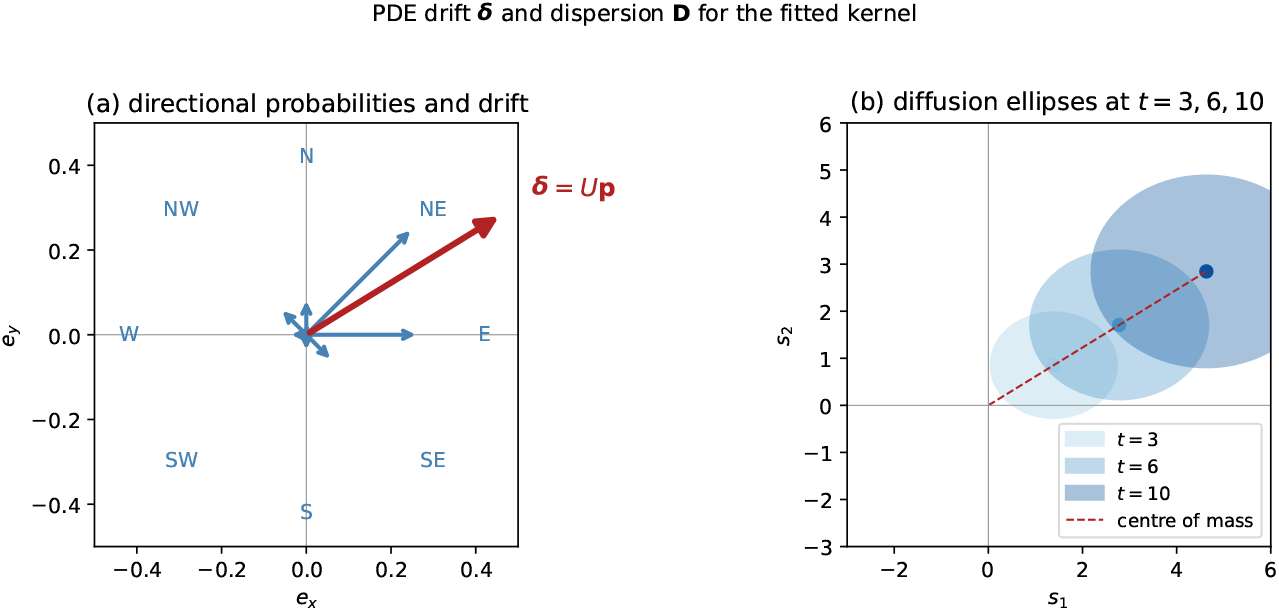
(a) Length-8 directional probability vector ***p***_true_ shown as blue arrows from the origin, with the resultant drift ***δ*** = *U* ***p*** in red. (b) Diffusion ellipses at *t* = 3, 6, 10 whose centres lie on the drift line ***δ****t* and whose semi-axes are 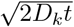.

## 7 Simulation Study

We simulate on a 25 × 25 grid with persistence *ϕ* = 0.85, long-distance dispersal *ψ* = 0.001, and anisotropic Dirichlet weights ***a***_true_ = (1, 4, 3, 1, 0.5, 0.2, 0.5, 1.0) ordered as (N, NE, E, SE, S, SW, W, NW). A single occupied seed is placed at the centre and the process is run for *T* = 20 steps. We then run 1500 MCMC iterations starting from *ϕ* = 0.5, *ψ* = 0.05, log ***a*** = **0**, with a Gaussian RW proposal of standard deviation 0.15 on log ***a***. Posterior means after a 500-iteration burn-in recover the true parameters within Monte Carlo error (*ϕ, ψ*, and ***p*** match the truth to two decimal places). Figure 4 shows snapshots of the simulated occupancy field, in which the NE-biased kernel ***p***_true_ drives an anisotropic front from the initial seed.

**Figure 4.**
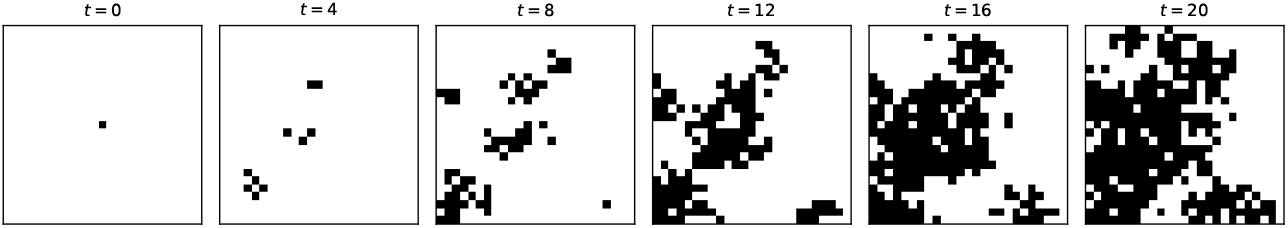
Stationary anisotropic SABM on a 25 × 25 grid with *ϕ* = 0.85, *ψ* = 0.001, and NE-biased kernel ***a***_true_ = (1, 4, 3, 1, 0.5, 0.2, 0.5, 1.0). Black cells are occupied. Starting from a single occupied centre cell at *t* = 0, the front advances most rapidly toward the northeast, consistent with the dominant directional probabilities *p*_NE_, *p*_E_.

For the PDE scaling check we run 5 000 independent Lagrangian random walks on a 41 × 41 grid starting at the centre, with kernel ***p***_true_, for 15 steps. The empirical agent density is compared to the explicit finite-difference solution of (13) with coefficients computed from (14). The root-mean-square error between the two fields is ≈ 4 × 10^−3^, and the centre of mass of the PDE solution coincides with the analytical drift (*δ*_1_*T, δ*_2_*T*) to within a fraction of a grid cell. This confirms numerically that the discrete kernel and the continuum PDE agree in the regime where the small-step approximation holds. Figure 5 contrasts the agent density and the PDE solution side by side at three snapshots: both fields advect to the northeast at the same rate and broaden at the same Gaussian width.

**Figure 5.**
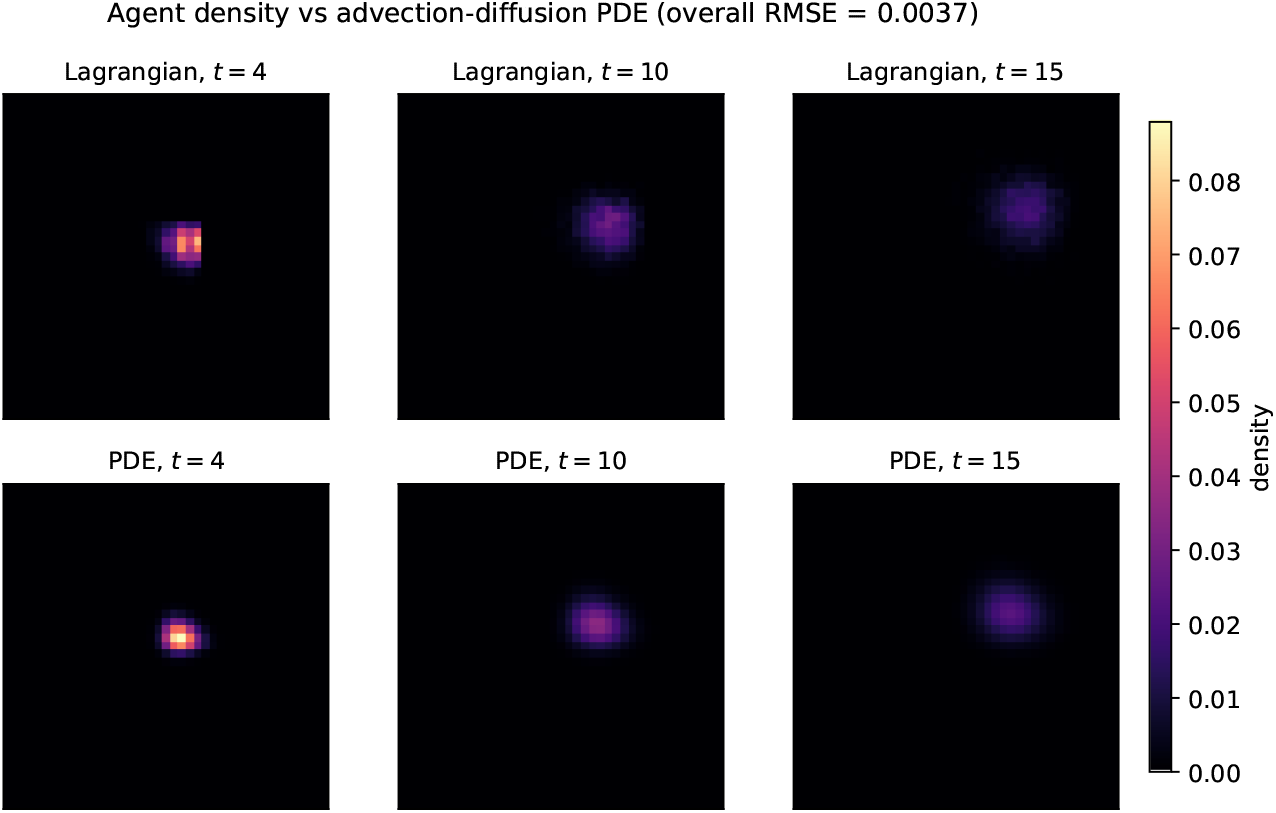
Lagrangian agent density (top row) from 5,000 random walks with kernel ***p***_true_, compared to the explicit finite-difference solution of the advection–diffusion PDE (13) with coefficients (***δ, D***) from (14) (bottom row). The two fields agree in shape, drift direction, and spread.

## 8 Application: Hudson’s Bay Lynx–Hare Series

We validate the SABM on the canonical Hudson’s Bay Company pelt record for the Canada lynx (*Lynx canadensis*) and snowshoe hare (*Lepus americanus*) over 1900–1920 (*n* = 21 years, counts in thousands of pelts; MacLulich, 1937; Elton and Nicholson, 1942). Each species is binarized at its median 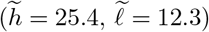, yielding

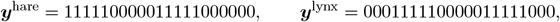

which exhibits the textbook out-of-phase predator–prey cycle with the lynx peak lagging the hare peak by roughly two years; the asymmetry of the cycle and its ~10-year period have been extensively analyzed (Stenseth et al., 1997; Bjørnstad and Grenfell, 2001). Figure 6 shows the underlying pelt counts with the median thresholds used for binarization.

**Figure 6.**
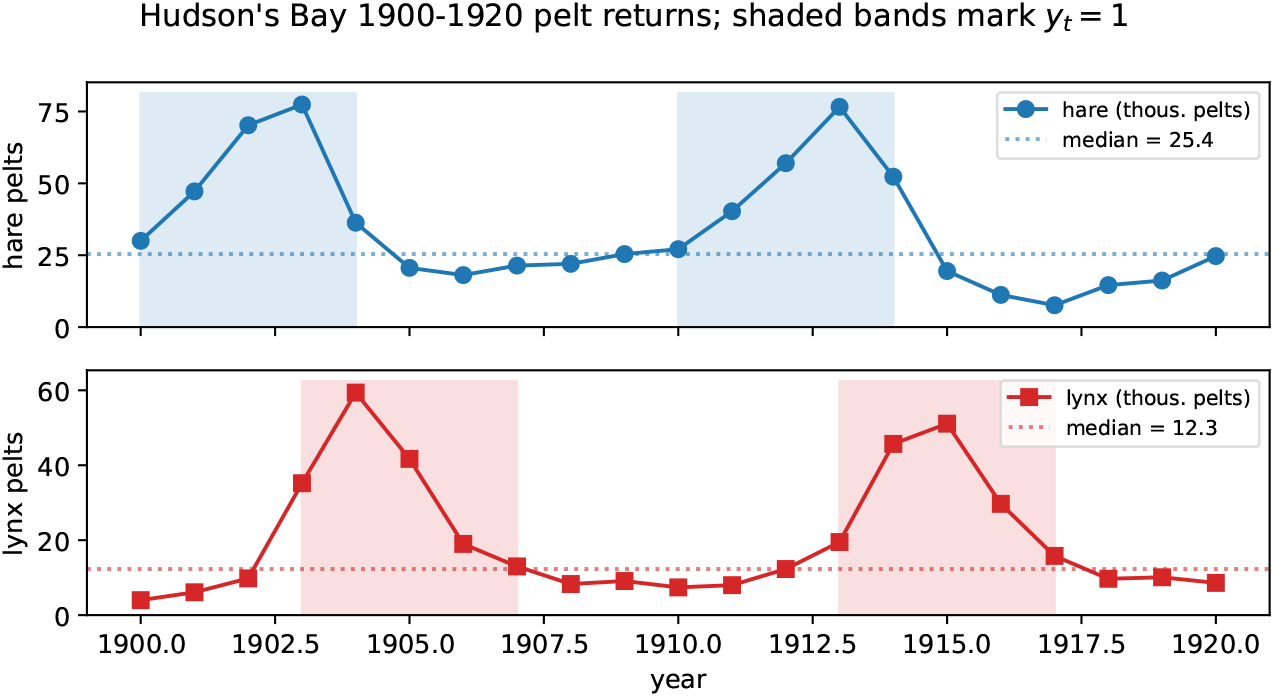
Hudson’s Bay Company hare and lynx pelt returns, 1900–1920 (thousands of pelts). Dotted lines mark the species-specific medians used as the binarization thresholds; shaded bands indicate years in which *y*_*t*_ = 1. The lynx peaks follow the hare peaks by about two years.

### Two-cell configuration

We place lynx in column 1 and hare in column 2 of a 1 × 2 grid, so the two species act as each other’s only on-grid neighbor. Of the eight directional entries of ***p*** only *p*_3_ (east: lynx → hare arrival) and *p*_7_ (west: hare → lynx arrival) are data-informed; the remaining six are sampled from their log-Gaussian prior. We ran the sampler in Algorithm 1 for 4,000 iterations and discarded the first 1,000 as burn-in; the Metropolis–Hastings acceptance for log ***a*** was 0.73.

**Table 1.** Posterior summaries for the two-cell SABM on the Hudson’s Bay hare/lynx series. The kernel asymmetry *p*_3_ ≫ *p*_7_ reflects the observed phase lag: hare booms are more often followed by lynx booms than the reverse.

| Parameter | Posterior mean | 95% CrI |
| --- | --- | --- |
| Persistence $\phi$ | 0.776 | [0.583, 0.918] |
| Long-distance $\psi$ | 0.200 | [0.029, 0.489] |
| $p_3$ (lynx → hare) | 0.246 | — |
| $p_7$ (hare → lynx) | 0.019 | — |

### Posterior predictive check

Forward-simulating 1,000 replicates from the posterior-mean parameters and the observed initial conditions, we compare both marginal occupancy and lag-1 transition counts to the data. Observed lynx occupancy is 0.48 vs. a predictive mean of 0.45 (95% interval [0.10, 0.81]); hare is 0.48 vs. 0.44 ([0.10, 0.90]). Lag-1 transition counts agree similarly: for lynx the four patterns 00*/*01*/*10*/*11 are observed as 8*/*2*/*2*/*8 versus predictive means 8.6*/*2.5*/*2.0*/*6.9; for hare 9*/*1*/*2*/*8 versus 9.9*/*1.3*/*1.9*/*6.9. All observed counts lie inside the 95% predictive intervals. Figure 7 summarizes the marginal posteriors and the predictive–observed comparison.

**Figure 7.**
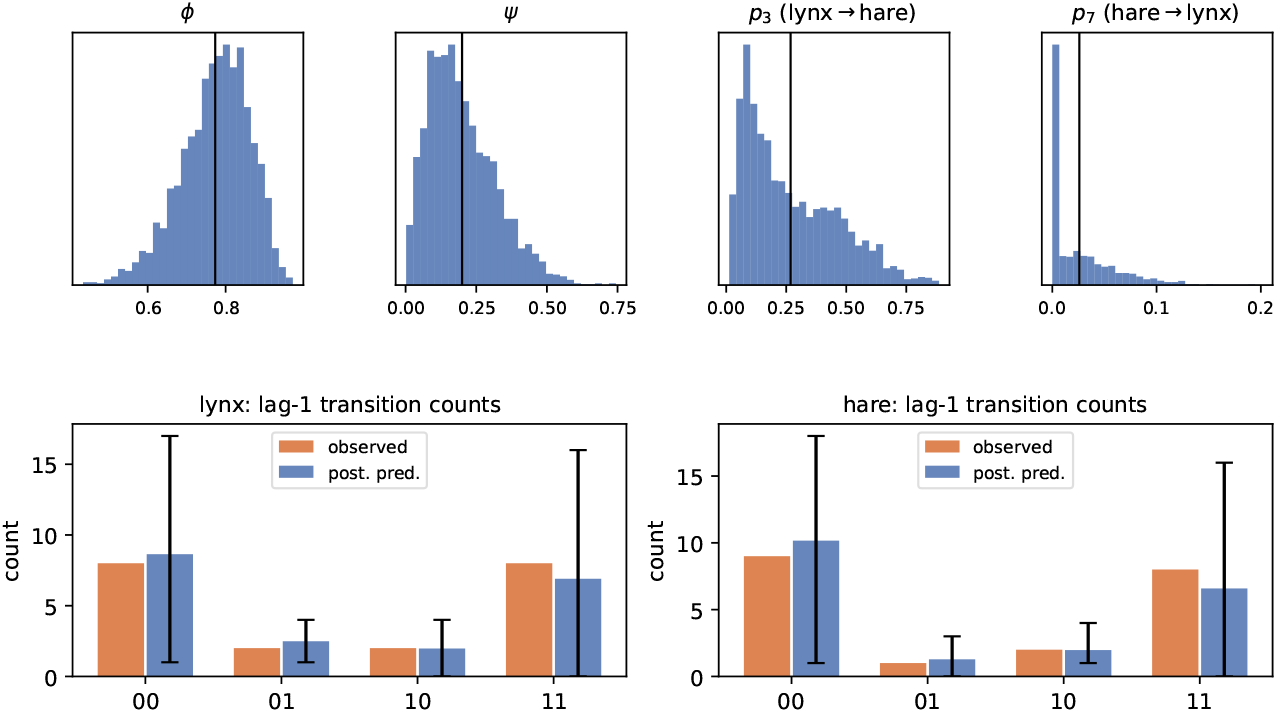
Top row: marginal posteriors of the two-cell SABM on the Hudson’s Bay series, for *ϕ, ψ*, and the data-informed directional entries *p*_3_, *p*_7_ (vertical line at the posterior mean). Bottom row: observed lag-1 transition counts (00*/*01*/*10*/*11) for lynx and hare (orange) against 1,000 posterior predictive replicates (blue, with 95% intervals). All observations sit inside the predictive intervals.

### One-step-ahead held-out accuracy

Refitting on the first 16 years (1900–1915) and scoring on 1916–1920 (ten species-year predictions) gives a Brier score of 0.116 and 9*/*10 correct point predictions at threshold 0.5. The single error is the 1918 lynx transition (1 → 0), which the model assigns probability 0.79 of remaining high; this is the year the cycle resets abruptly, and the most difficult prediction in the series. Figure 8 overlays the observed binarized series with the one-step-ahead predictive probabilities for the held-out window, together with 95% credible bands obtained from the posterior draws.

**Figure 8.**
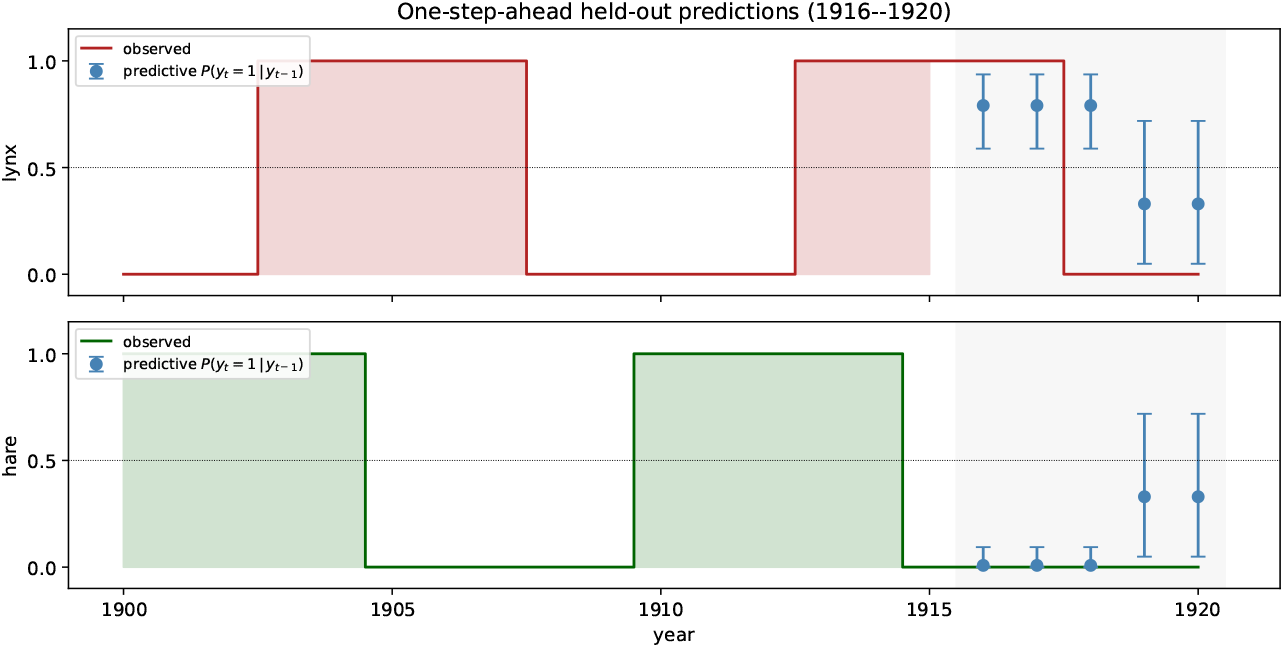
Held-out one-step-ahead predictions for the two-cell lynx–hare SABM. Solid step traces are the observed binarized series (*y*_*t*_ = 1 if pelt count exceeds the species median); shaded panels indicate the training window 1900–1915. Blue points with whiskers are the posterior-mean pre-dictive probabilities *P* (*y*_*t*_ = 1 | *y*_*t*−1_) for 1916–1920 and the corresponding 95% credible intervals. Dotted line at 0.5 marks the decision threshold.

### Interpretation

The calibration of the predictive intervals and the held-out accuracy support the SABM as a faithful generative description of binary occupancy dynamics on real ecological data. The two-cell configuration recovers the directional asymmetry of the predator–prey lag without the model having any predator–prey machinery built in: the spread mechanism alone reproduces lag-1 statistics within Monte Carlo error. Spatially resolved Hudson’s Bay returns by district would let the full eight-direction kernel be estimated and is a natural next step.

## 9 Discussion

The SABM places mechanistic spread on a rigorous statistical footing. Three ingredients combine to make it tractable: disjoint partitioning of *θ* into persistence, neighborhood, and long-distance regimes; a Dirichlet kernel that admits a log-Gaussian prior on its concentration; and a gradient-driven nonstationary extension that ties dispersal to a Gaussian-process suitability surface. The exact *Beta* full-conditionals for *ϕ* and *ψ* sharply reduce the MCMC variance for those parameters, the Lynx–Hare application confirms that this component recovers cycle lengths consistent with classical ecological theory, and the PDE limit gives a direct bridge to deterministic spread models for sanity checking and for extrapolation beyond the data.

Several extensions are natural: vectorized site-level covariates in *X*; a fully Bayesian treatment of *σ*_*α*_, *θ*_*α*_ via slice or Metropolis–Hastings; multi-state occupancy (e.g. susceptible, infected, recovered) by enlarging the alphabet of *y*_*i,t*_; and time-varying *ϕ, ψ* when external drivers (weather, control measures) are present.

## Data and Code Availability

The Hudson’s Bay Company lynx–hare pelt records used in the application are public-domain historical data (MacLulich, 1937; Elton and Nicholson, 1942) and are included verbatim in the project repository at `data/lynx hare.csv`. All R source files implementing the SABM, the MCMC sampler, the PDE scaling check, and the lynx–hare validation are available in the same repository (link to be provided on acceptance) and are reproduced in full in the Supplementary Material. The analyses in this paper run end-to-end via `Rscript demo.R` (simulation study) and `Rscript validate_lynx_hare.R` (data application).

## Author Contributions

X.P. conceived the model, implemented the software, performed the analyses, and wrote the manuscript.

## Acknowledgments

The author thanks the Penn State community for useful discussions on hierarchical Bayesian spatio-temporal models.

---

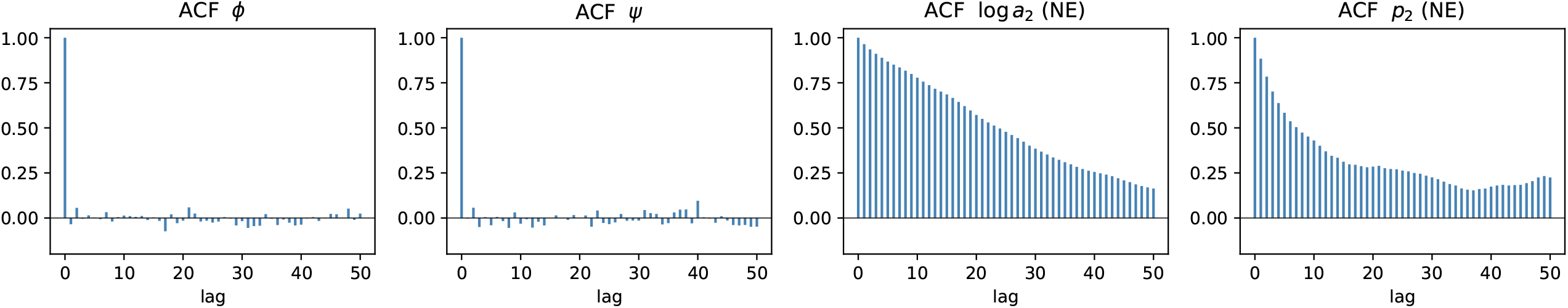

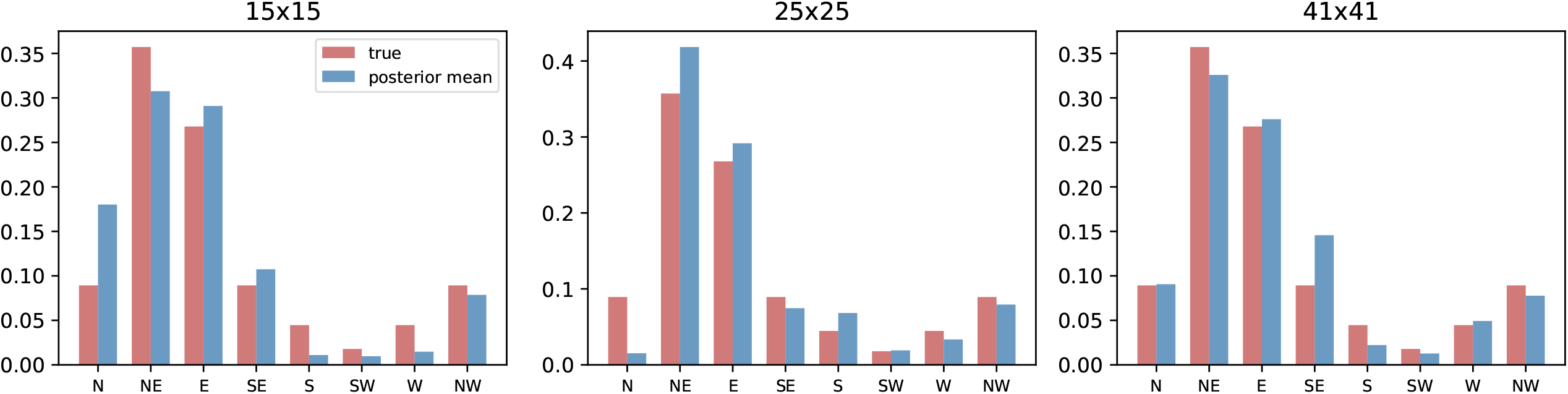

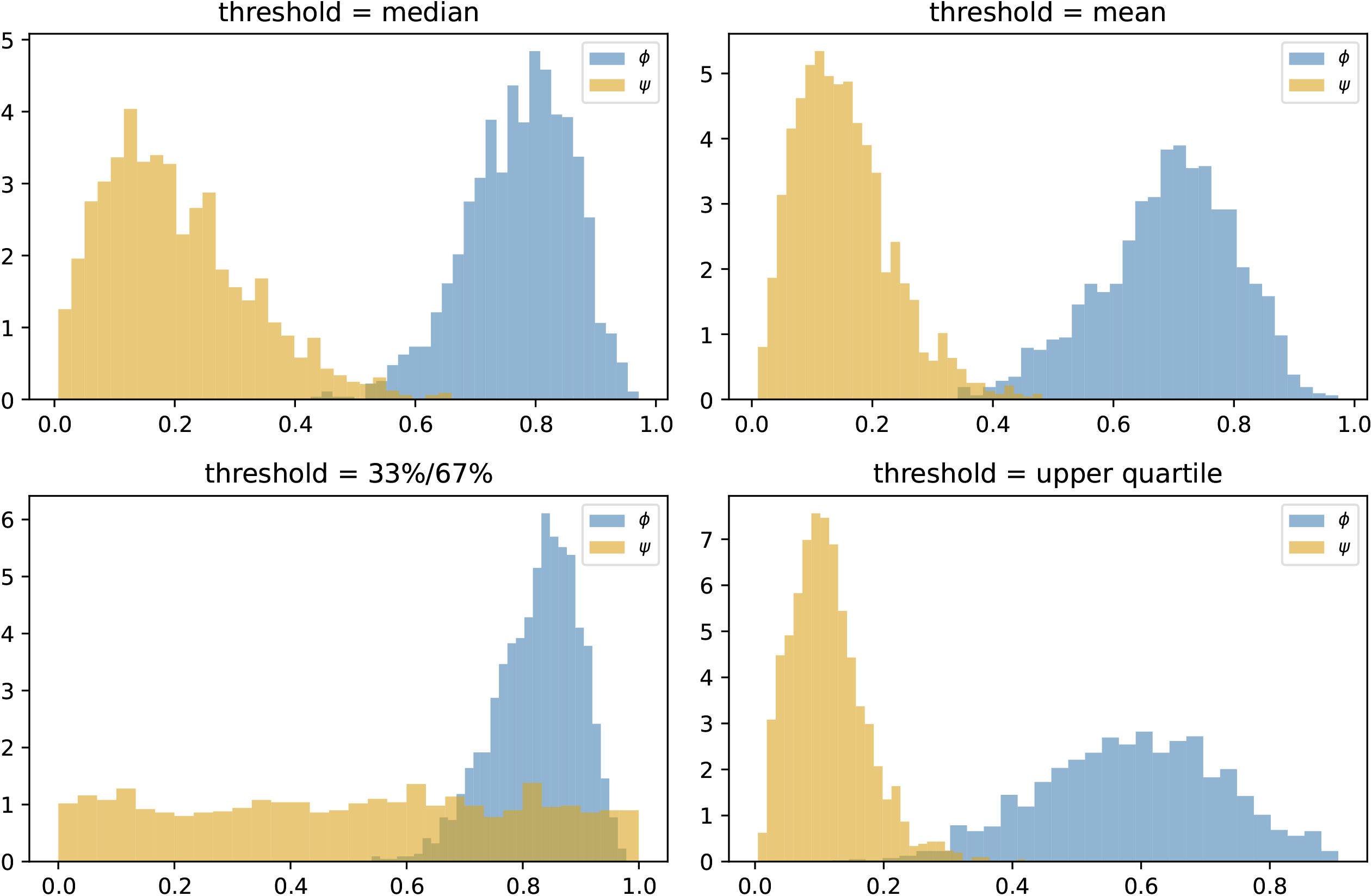

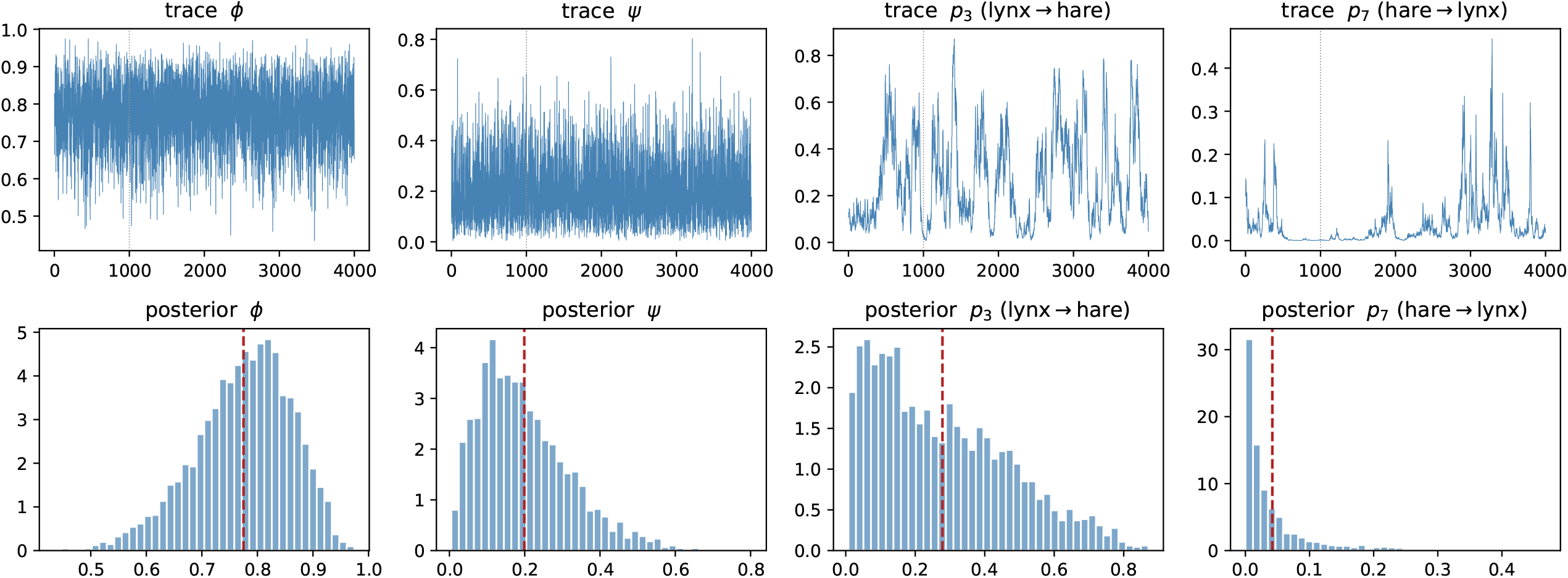

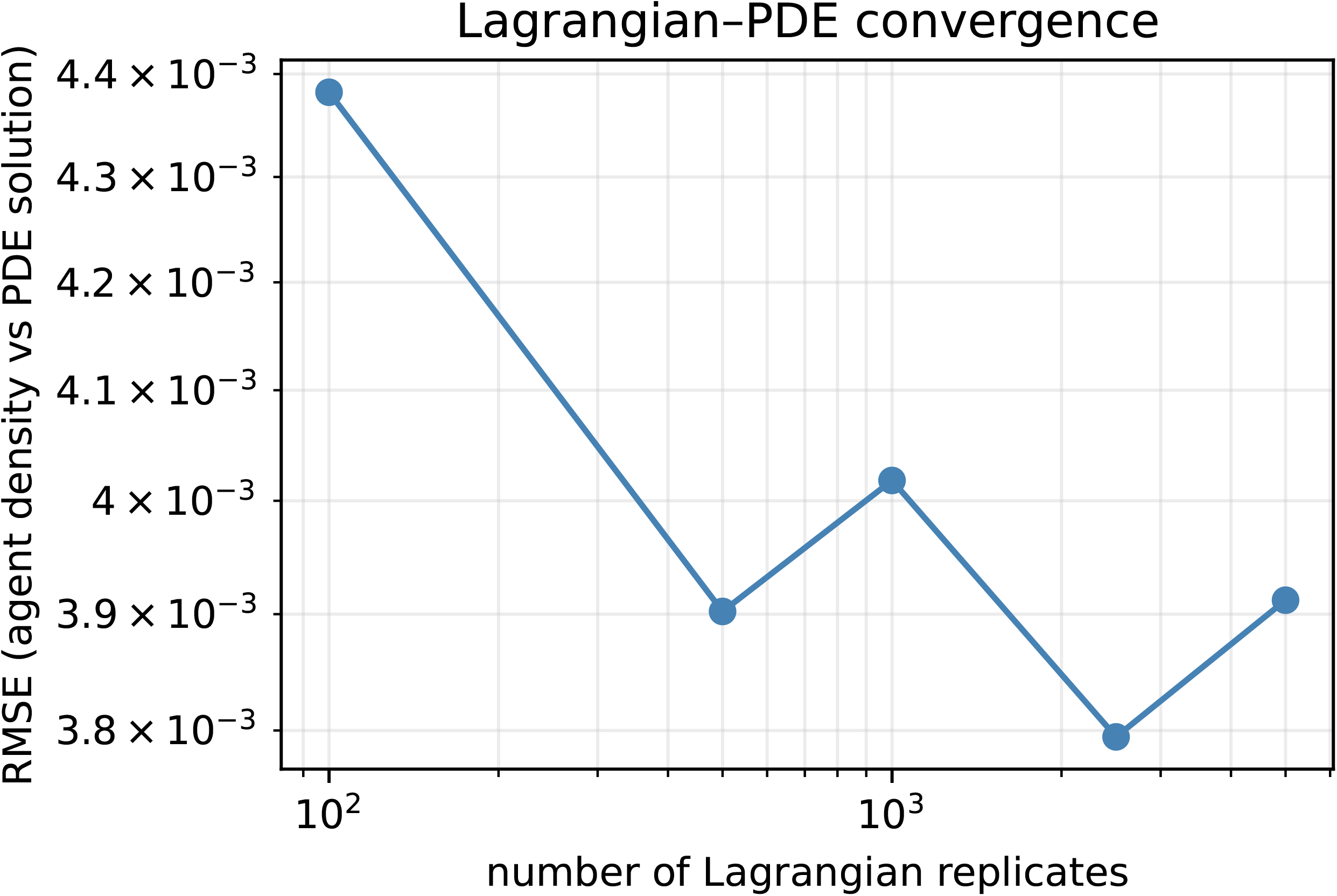

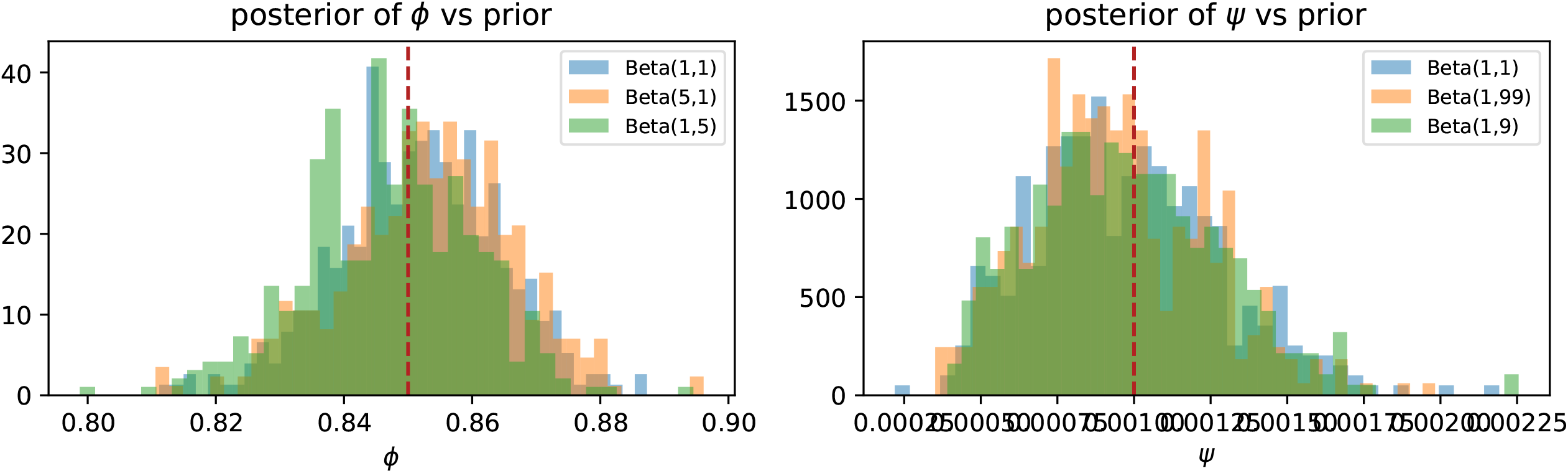

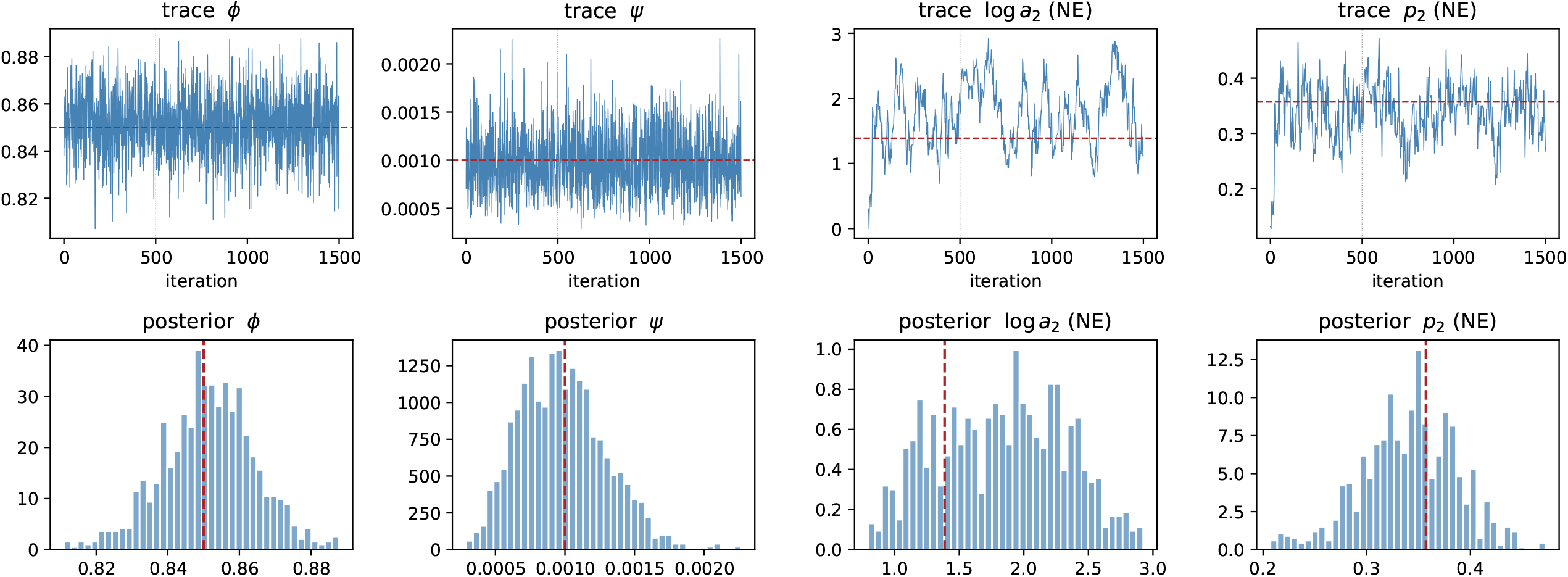

