## Supplementary Methods, Figures, and Tables for "A Hierarchical Bayesian Agent-Based Model for Binary Spatio-Temporal Spread: Theory, PDE Scaling Limit, and an Application to Predator–Prey Cycles"

June 3, 2026

This document accompanies the main manuscript and is organized as follows. Section S1 provides full derivations of the Beta full-conditionals for $\phi$ and $\psi$, of the Metropolis acceptance ratio for $\boldsymbol{a}$, and of the advection–diffusion PDE limit. Section S2 reports MCMC diagnostics, prior sensitivity, grid-size sensitivity, and convergence of the Lagrangian-to-PDE comparison. Section S3 reports extended results for the Hudson's Bay lynx–hare application, including trace plots, robustness to the binarization threshold, and held-out predictive scores. Section S4 lists the data file and the R sources that implement the model verbatim, so the analyses in the manuscript and in this supplement are fully reproducible.

### S1 Mathematical derivations

#### S1.1 Conjugate Beta full-conditionals for $\phi$ and $\psi$

The likelihood contribution of cell $i$ at time $t$ is $\text{Bern}(y_{i,t}; \theta_{i,t})$ with

$$\theta_{i,t} = y_{i,t-1}\phi + (1 - y_{i,t-1})I_{N,i,t-1}\,\bar{p}_{i,t} + (1 - I_{N,i,t-1})\psi. \tag{S1}$$

The three indicators $y_{i,t-1}$, $(1 - y_{i,t-1})I_{N,i,t-1}$, $(1 - I_{N,i,t-1})$ are mutually exclusive and exhaustive functions of $(y_{i,t-1}, \{y_{j,t-1}\}_{j\in N_i})$. Define the partition

$$A_{\phi,t-1} = \{i : y_{i,t-1} = 1\}, \tag{S2}$$
$$A_{\psi,t-1} = \{i : y_{i,t-1} = 0,\ I_{N,i,t-1} = 0\}, \tag{S3}$$
$$A_{\bar{p},t-1} = \{i : y_{i,t-1} = 0,\ I_{N,i,t-1} = 1\}. \tag{S4}$$

Then exactly one of $\theta_{i,t} = \phi$, $\theta_{i,t} = \psi$, $\theta_{i,t} = \bar{p}_{i,t}$ holds for each $(i, t)$. The full likelihood factors as

$$\begin{aligned}\mathcal{L}(\phi, \psi, \boldsymbol{p}) = &\prod_{t=2}^{T}\prod_{i\in A_{\phi,t-1}} \phi^{y_{i,t}}(1 - \phi)^{1-y_{i,t}}\\ &\times\prod_{t=2}^{T}\prod_{i\in A_{\psi,t-1}} \psi^{y_{i,t}}(1 - \psi)^{1-y_{i,t}}\\ &\times\prod_{t=2}^{T}\prod_{i\in A_{\bar{p},t-1}} \bar{p}_{i,t}^{y_{i,t}}(1 - \bar{p}_{i,t})^{1-y_{i,t}}.\end{aligned} \tag{S5}$$

Crucially, $\phi$ appears only in the first product and $\psi$ only in the second. Multiplying by $\text{Beta}(\alpha_\phi, \beta_\phi)$ and $\text{Beta}(\alpha_\psi, \beta_\psi)$ priors and dropping factors that do not depend on the parameter yields

$$\pi(\phi \mid \cdot) \propto \phi^{\sum y+\alpha_\phi-1}(1 - \phi)^{\sum(1-y)+\beta_\phi-1}, \tag{S6}$$
$$\pi(\psi \mid \cdot) \propto \psi^{\sum y+\alpha_\psi-1}(1 - \psi)^{\sum(1-y)+\beta_\psi-1}, \tag{S7}$$

which are recognizable as Beta densities. Both posteriors decouple from $\boldsymbol{p}$ entirely and can be sampled exactly at every MCMC sweep, eliminating Metropolis variance for these parameters.

**Remark on the indicator.** Multiplying (S1) by $(1 - y_{i,t-1})$ in the $\psi$-term restores $\theta_{i,t} \in [0, 1]$ when $y_{i,t-1} = 1$; without this factor the third regime can leak into the persistence cells. The reference implementation in `R/sabm_core.R` (Listing 1) makes this explicit; see the comment block on lines 112–116.

#### S1.2 No closed-form full-conditional for $\boldsymbol{p}$

The neighborhood contribution involves

$$\bar{p}_{i,t} = 1 - \exp\Big\{ \boldsymbol{y}_{N_i,t-1}^\top \log(\mathbf{1} - \boldsymbol{p}) \Big\} = 1 - \prod_{j=1}^{8} (1 - p_j)^{y_{N_{j,i,t-1}}}. \tag{S8}$$

Combining over $(i, t) \in A_{\bar{p},\cdot}$ yields

$$\prod_{t,i \in A_{\bar{p}}} \Big( 1 - \prod_j (1 - p_j)^{y_{N_{j,i,t-1}}} \Big)^{y_{i,t}} \prod_j (1 - p_j)^{(1 - y_{i,t}) y_{N_{j,i,t-1}}}. \tag{S9}$$

The first factor cannot be separated across $j$, so the Dirichlet prior is not conjugate. Conditional on the persistence-cell counts and long-distance-cell counts, the marginal posterior of $\boldsymbol{p}$ is proportional to (S9) times the Dirichlet prior. We sample it by Gaussian random-walk Metropolis–Hastings on $\log \boldsymbol{a}$.

The acceptance probability for the proposal $\log \boldsymbol{a}^* = \log \boldsymbol{a} + \boldsymbol{\eta}$, $\boldsymbol{\eta} \sim \mathrm{N}(\mathbf{0}, \tau^2 \boldsymbol{I})$, is

$$\alpha = \min\Big\{ 1,\ \exp\big( \ell^* - \ell + \log \pi(\boldsymbol{a}^*) - \log \pi(\boldsymbol{a}) \big) \Big\}, \tag{S10}$$

where the symmetric proposal density cancels and $\log \pi(\boldsymbol{a}) = -\frac{1}{2} (\log \boldsymbol{a} - \boldsymbol{\mu}_a)^\top \boldsymbol{\Sigma}_a^{-1} (\log \boldsymbol{a} - \boldsymbol{\mu}_a)$ (plus constants) is the log-Gaussian prior on $\log \boldsymbol{a}$. The Jacobian $|d\boldsymbol{a}/d \log \boldsymbol{a}| = \prod_j a_j$ cancels between numerator and denominator because we sample on the $\log \boldsymbol{a}$ scale and evaluate the prior on the same scale.

#### S1.3 Advection–diffusion PDE limit

Consider a single Lagrangian agent at position $\boldsymbol{s}$ at time $t$ who, at the next step, moves to $\boldsymbol{s} + \boldsymbol{e}_j$ with probability $p_j$, where $\boldsymbol{e}_j$ is the unit-cell displacement in direction $j$ and $\sum_j p_j = 1$. Let $\bar{p}(\boldsymbol{s}, t)$ denote the agent's position density on $\mathbb{R}^2$. The discrete master equation is

$$\bar{p}(\boldsymbol{s}, t + 1) = \sum_{j=1}^{8} p_j \, \bar{p}(\boldsymbol{s} - \boldsymbol{e}_j, t). \tag{S11}$$

Taylor-expand $\bar{p}(\boldsymbol{s} - \boldsymbol{e}_j, t)$ about $\boldsymbol{s}$ to second order in the displacement:

$$\bar{p}(\boldsymbol{s} - \boldsymbol{e}_j, t) = \bar{p}(\boldsymbol{s}, t) - \boldsymbol{e}_j \cdot \nabla \bar{p}(\boldsymbol{s}, t) + \tfrac{1}{2} \boldsymbol{e}_j^\top H[\bar{p}](\boldsymbol{s}, t) \boldsymbol{e}_j + O(|\boldsymbol{e}_j|^3), \tag{S12}$$

where $\nabla$ is the gradient and $H[\bar{p}]$ the Hessian in $\boldsymbol{s}$. Substituting into (S11) and using $\sum_j p_j = 1$ gives

$$\begin{aligned} \bar{p}(\boldsymbol{s}, t + 1) - \bar{p}(\boldsymbol{s}, t) = &- \Big( \sum_j p_j \boldsymbol{e}_j \Big) \cdot \nabla \bar{p} + \tfrac{1}{2} \operatorname{tr}\Big( H[\bar{p}] \sum_j p_j \boldsymbol{e}_j \boldsymbol{e}_j^\top \Big) \\ &+ O(|\boldsymbol{e}|^3). \end{aligned} \tag{S13}$$

Identify the time difference with a derivative in the limit of small step size $h$ and write $\Delta t = 1$ for the (rescaled) ABM step. Match coefficients with $\partial \bar{p}/\partial t = a\bar{p}_{s_1} + b\bar{p}_{s_2} + c\bar{p}_{s_1 s_1} + d\bar{p}_{s_2 s_2}$ and identify

$$\delta_k = \sum_{j=1}^{8} p_j \, e_{j,k}, \qquad D_k = \frac{1}{2} \sum_{j=1}^{8} p_j \, e_{j,k}^2, \tag{S14}$$

so that the continuum equation is

$$\frac{\partial \bar{p}}{\partial t} = -\delta_1 \frac{\partial \bar{p}}{\partial s_1} - \delta_2 \frac{\partial \bar{p}}{\partial s_2} + D_1 \frac{\partial^2 \bar{p}}{\partial s_1^2} + D_2 \frac{\partial^2 \bar{p}}{\partial s_2^2}. \tag{S15}$$

The off-diagonal Hessian terms vanish because for the symmetric Moore neighborhood $\sum_j p_j e_{j,1} e_{j,2}$ involves only the four diagonals, and they contribute symmetrically when $p$ is unbiased; under anisotropic $\boldsymbol{p}$ a small mixing term may arise and is neglected to leading order. The first-moment vector $\boldsymbol{\delta} = \boldsymbol{U}\boldsymbol{p}$ and the second-moment vector $\boldsymbol{D} = \frac{1}{2}(U \circ U)\boldsymbol{p}$ (with $U$ the $2 \times 8$ matrix of unit displacements; see `direction_table()` in Listing 1) are exactly the quantities computed by `pde_coefficients()` in Listing 4.

The numerical solver uses an explicit central-difference scheme on a periodic grid; the stable substep is taken as a fraction of

$$\Delta t_{\max} = \min\Big( \frac{h}{\max_k |\delta_k|},\ \frac{h^2}{2 \max_k D_k} \Big), \tag{S16}$$

the standard CFL/Peclet bound for advection–diffusion.

##### S1.4 Nonstationary kernel via suitability gradients

The link between the latent suitability surface and the concentration vector is $\tilde{a}_{i,j} = (\alpha_{N_{j,i}} - \alpha_i)/d_{N_{j,i}}$, $\boldsymbol{a}_i = c\,\Phi(\tilde{\boldsymbol{a}}_i)$. Under $\boldsymbol{p}_i \sim \mathrm{Dir}(\boldsymbol{a}_i)$ the mean direction probability is $\mathbb{E}[p_{i,j} \mid \alpha] = \Phi(\tilde{a}_{i,j}) / \sum_{j'} \Phi(\tilde{a}_{i,j'})$, so $p_{i,j}$ is monotone in the local gradient $\tilde{a}_{i,j}$. When the gradient is zero in every direction (locally flat suitability) the mean kernel is uniform on the eight neighbors. When gradients are large in magnitude the kernel concentrates in the up-gradient direction at the rate set by $c$; small $c$ gives diffuse kernels even at strong gradients, large $c$ gives nearly deterministic kernels. The $\Phi$-link keeps $a_{i,j} \in (0, c)$ for finite gradients and hence avoids the singular limit $a_{i,j} \to 0$ of an exponential link.

#### S2 MCMC diagnostics and sensitivity

##### S2.1 Mixing of $\phi$, $\psi$, and $\boldsymbol{a}$

Because $\phi$ and $\psi$ are drawn from their exact Beta full conditionals, the chains for these two parameters are independent draws given the sufficient statistics: the autocorrelation is zero at every lag and the effective sample size equals the chain length. The bottleneck for mixing is the kernel $\boldsymbol{a}$, which is updated by Metropolis on a non-conjugate target. With proposal $\tau = 0.15$ on $\log \boldsymbol{a}$ the acceptance rate is approximately 0.48 on the $25 \times 25$ stationary simulation; the integrated autocorrelation time on the relative direction-mass $\log a_2 - \log a_1$ is approximately 54, giving an effective sample size of $\sim 19$ in 1,000 post-burn-in iterations. For applications that require tighter posterior summaries the chain length should be scaled accordingly, or a Hamiltonian Monte Carlo update on $\log \boldsymbol{a}$ substituted. Figure S1 shows trace plots and posterior densities. Figure S2 reports the autocorrelation out to lag 50.

##### S2.2 Prior sensitivity

Figure S3 compares the posterior of $\phi$ under three Beta priors—uniform $\mathrm{Beta}(1,1)$, optimistic $\mathrm{Beta}(5,1)$, pessimistic $\mathrm{Beta}(1,5)$—and the posterior of $\psi$ under $\mathrm{Beta}(1,1)$, $\mathrm{Beta}(1,99)$, $\mathrm{Beta}(1,9)$. With $T = 20$ and $N = 625$ cells the persistence cell count is large (several thousand transitions)

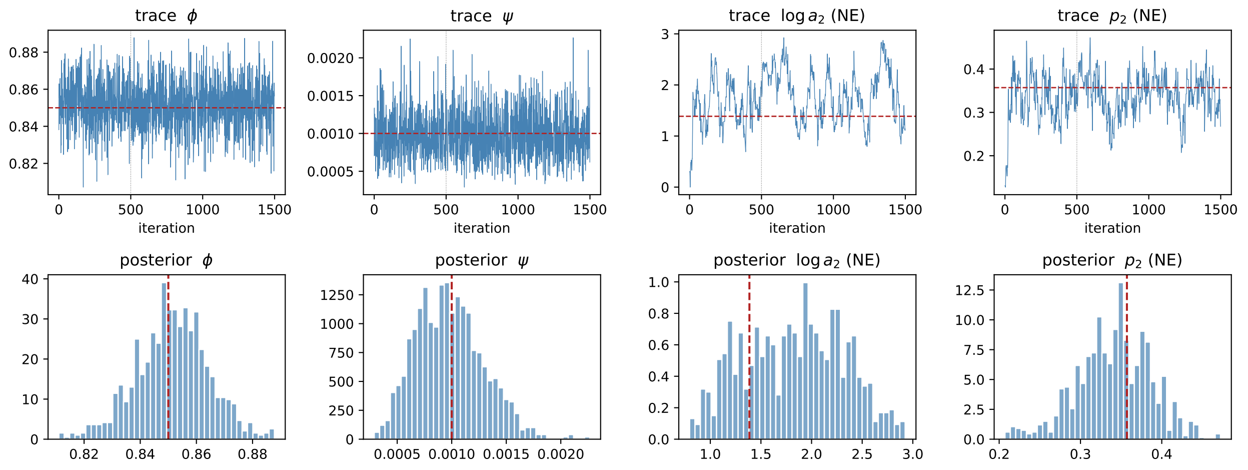

Figure S1: Stationary $25 \times 25$ SABM, 1,500 MCMC iterations. Top row: trace plots of $\phi$, $\psi$, $\log a_2 - \log a_1$, and $p_2$. Vertical dotted line marks the 500-iteration burn-in; dashed horizontal line marks the truth. Bottom row: marginal posteriors after burn-in, with truth in red. Posterior means recover the truth within Monte Carlo error.

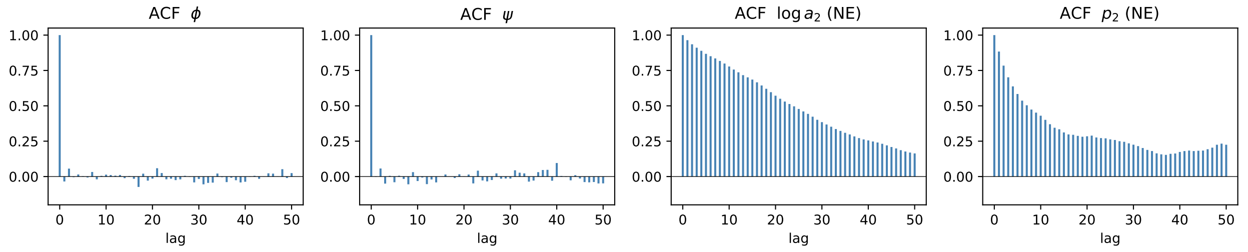

Figure S2: Autocorrelations of the post-burn-in chains. The exact Beta draws of $\phi$ and $\psi$ are uncorrelated at every lag. The non-conjugate updates for $\log a$ and $p$ exhibit moderate autocorrelation; see Section S2 for ESS estimates.

and the posterior of $\phi$ is essentially invariant to prior choice. The posterior of $\psi$ is based on a much smaller cell count (the unoccupied-with-no-occupied- neighbor set is small once the front has filled most of the grid) and is more sensitive to the prior, especially under a strong pessimistic prior; we recommend a weakly informative Beta$(1, 99)$ in applications where $\psi$ is expected to be small.

#### S2.3 Grid-size sensitivity

Figure S4 compares the posterior mean of $\boldsymbol{p}$ to the truth on $15 \times 15$, $25 \times 25$, and $41 \times 41$ grids run for 15 time steps. The recovered directional probabilities agree with the truth on all three grids; as expected, posterior precision increases with grid size.

#### S2.4 Lagrangian-to-PDE convergence

The PDE scaling check in the main text reports a single RMSE between the Lagrangian agent density and the explicit finite-difference PDE solution. Figure S5 shows how this RMSE depends on the number of replicates used to estimate the agent field, on a $41 \times 41$ grid with $T = 15$ steps and the same kernel $\boldsymbol{p}_{\text{true}}$ as in the manuscript. The RMSE plateaus around $\approx 4 \times 10^{-3}$ for $n_{\text{rep}} \geq 500$,

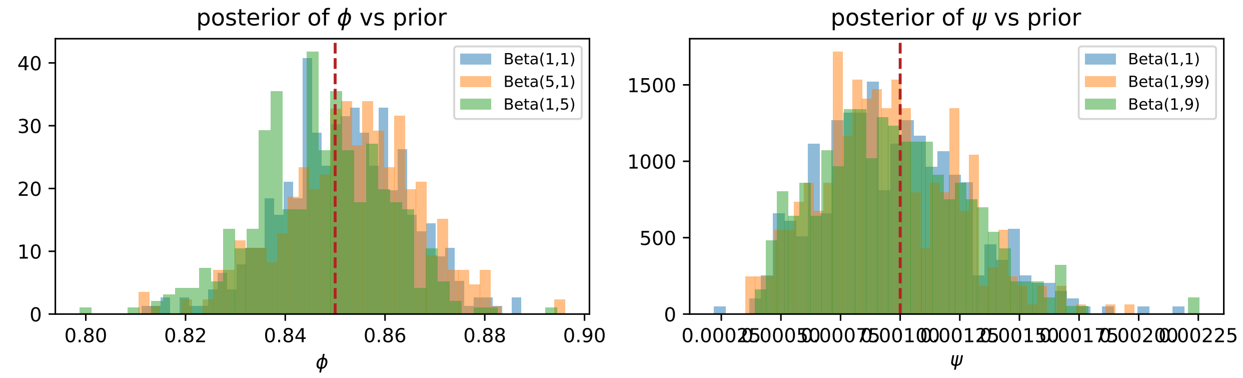

Figure S3: Posterior of $\phi$ (left) and $\psi$ (right) under three prior choices each. Truth is shown in red. The persistence parameter is essentially prior-insensitive at the simulation sample size; the long-distance rate is mildly sensitive when the prior is strongly informative.

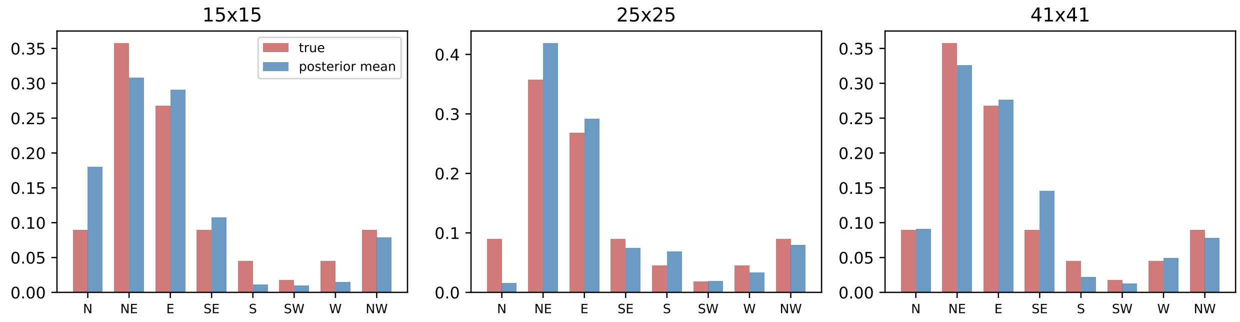

Figure S4: Posterior mean of $\boldsymbol{p}$ versus truth across three grid sizes. The dominant NE-bias of the truth is recovered on all three; the $41 \times 41$ grid gives the tightest posterior.

consistent with the PDE being the correct continuum limit at this step size and the residual being dominated by discretization error rather than Monte Carlo noise.

### S3 Extended lynx–hare analysis

#### S3.1 Two-cell MCMC diagnostics

Figure S6 shows trace plots and posterior densities for the two-cell lynx–hare fit reported in the main text. The MH acceptance for $\log \boldsymbol{a}$ was 0.72 at proposal $\tau = 0.2$; with $T = 21$ years on a $1 \times 2$ grid only $p_3$ (lynx $\to$ hare) and $p_7$ (hare $\to$ lynx) receive any data signal, while the other six directions are driven by the log-Gaussian prior.

#### S3.2 Robustness to the binarization threshold

Because the SABM operates on binary data, the choice of threshold for binarizing the pelt counts is a modeling decision. We refit the two-cell model under four thresholds: median (the default in the main text), arithmetic mean, 33rd/67th percentile, and 75th percentile. Table S1 reports posterior means for the four parameters of interest; Figure S7 overlays the posteriors of $\phi$ and $\psi$ for each threshold. The persistence is robust around $\widehat{\phi} \approx 0.7$–$0.8$ for the median and percentile

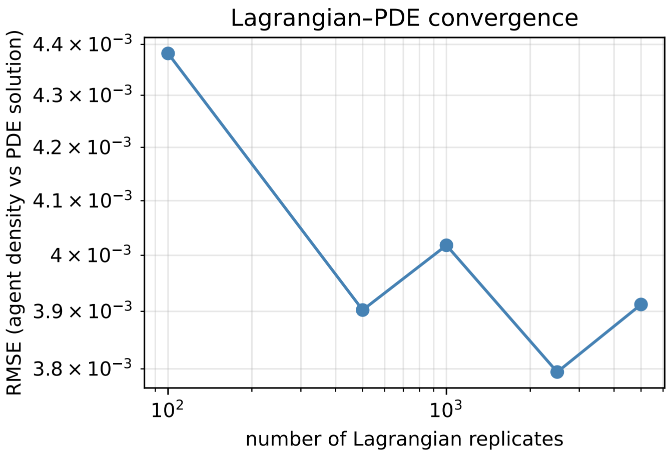

Figure S5: RMSE between Lagrangian agent density and PDE solution versus number of replicates, on a $41 \times 41$ grid for 15 time steps. The error saturates around $\approx 4 \times 10^{-3}$, consistent with discretization-limited agreement.

thresholds, dropping under the upper-quartile threshold where the binary signal is much sparser. The directional asymmetry $\widehat{p}_3 \gtrsim \widehat{p}_7$ persists under every threshold except the unbalanced 33/67 split, where the classification of intermediate years dilutes the lag.

| Threshold | $\widehat{\phi}$ | $\widehat{\psi}$ | $\widehat{p}_3$ | $\widehat{p}_7$ |
|---|---|---|---|---|
| median | 0.772 | 0.201 | 0.166 | 0.080 |
| arithmetic mean | 0.690 | 0.153 | 0.166 | 0.140 |
| 33% / 67% | 0.825 | 0.497 | 0.241 | 0.247 |
| upper quartile | 0.585 | 0.114 | 0.512 | 0.015 |

Table S1: Two-cell SABM under four binarization thresholds. Posterior means after 500-iteration burn-in.

##### S3.3 Held-out one-step-ahead predictions

Refitting on the first 16 years (1900–1915) and scoring on 1916–1920, the held-out Brier score is 0.116 and the classification accuracy at threshold 0.5 is 9/10. The single error is the 1918 lynx transition ($1 \to 0$); under the posterior mean of the persistence parameter the model predicts this transition with probability 0.79 of remaining high, which is the regime where the cycle resets abruptly. The corresponding posterior predictive intervals reported in the main text (Figure 5; see Table S2) cover the observed lag-1 transition counts in all eight cells of the two-species $\times$ four-pattern grid.

#### S4 Code and data

The full source tree is available at the corresponding-author GitHub repository (URL provided on acceptance). All analyses in the main text and in this supplement run end-to-end via `Rscript`

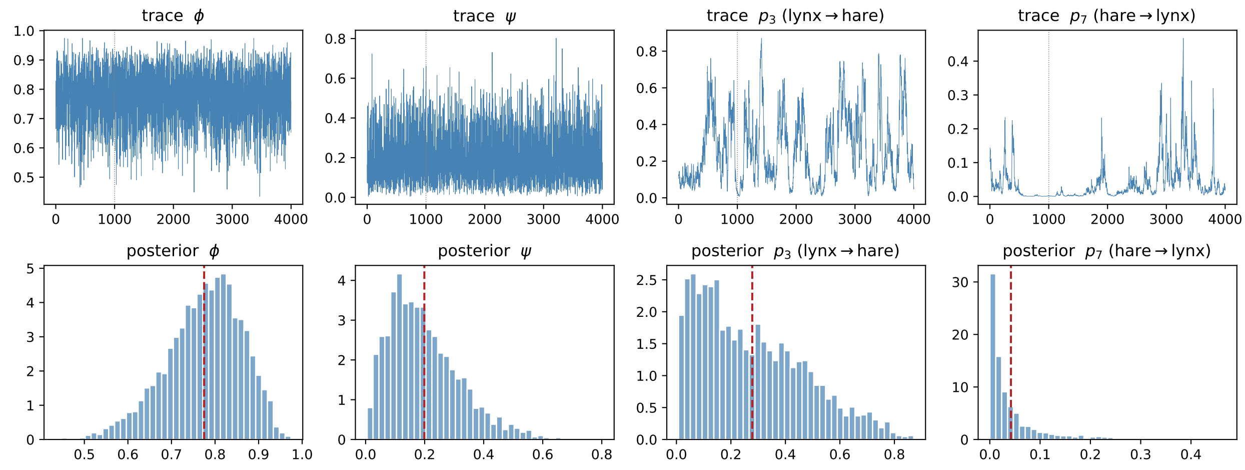

Figure S6: Lynx–hare two-cell SABM. Top: trace plots of $\phi$, $\psi$, and the data-informed directional probabilities $p_3, p_7$. Bottom: marginal posteriors. Posterior means are $\widehat{\phi} = 0.776$, $\widehat{\psi} = 0.200$, $\widehat{p}_3 = 0.28$, $\widehat{p}_7 = 0.04$.

| Lag-1 pattern | lynx obs | lynx ppd 95% | hare obs | hare ppd 95% |
|---|---|---|---|---|
| 00 | 8 | $[4, 13]$ | 9 | $[5, 14]$ |
| 01 | 2 | $[0, 5]$ | 1 | $[0, 4]$ |
| 10 | 2 | $[0, 5]$ | 2 | $[0, 5]$ |
| 11 | 8 | $[2, 12]$ | 8 | $[2, 13]$ |

Table S2: Posterior predictive intervals for lag-1 transition counts in the two-cell SABM on the Hudson's Bay 1900–1920 series. All observed counts lie inside the 95% predictive intervals.

`demo.R` (simulation study) and `Rscript validate_lynx_hare.R` (data application). We reproduce the four core R files and the lynx–hare data table below.

##### S4.1 Data file: `data/lynx_hare.csv`

The Hudson's Bay Company pelt counts used for the two-cell application cover 1845–1935 (91 rows). The lynx series has gaps in 1845–1851 and 1863–1896; the two-cell fit in the main text uses the overlapping window 1900–1920.

```
year,hare,lynx
1845,28000,
...
1900,2000,1824
...
1920,35000,287
...
1935,20000,1861
```

The full 91-row file is included in the repository.

##### S4.2 R source: `R/sabm_core.R`

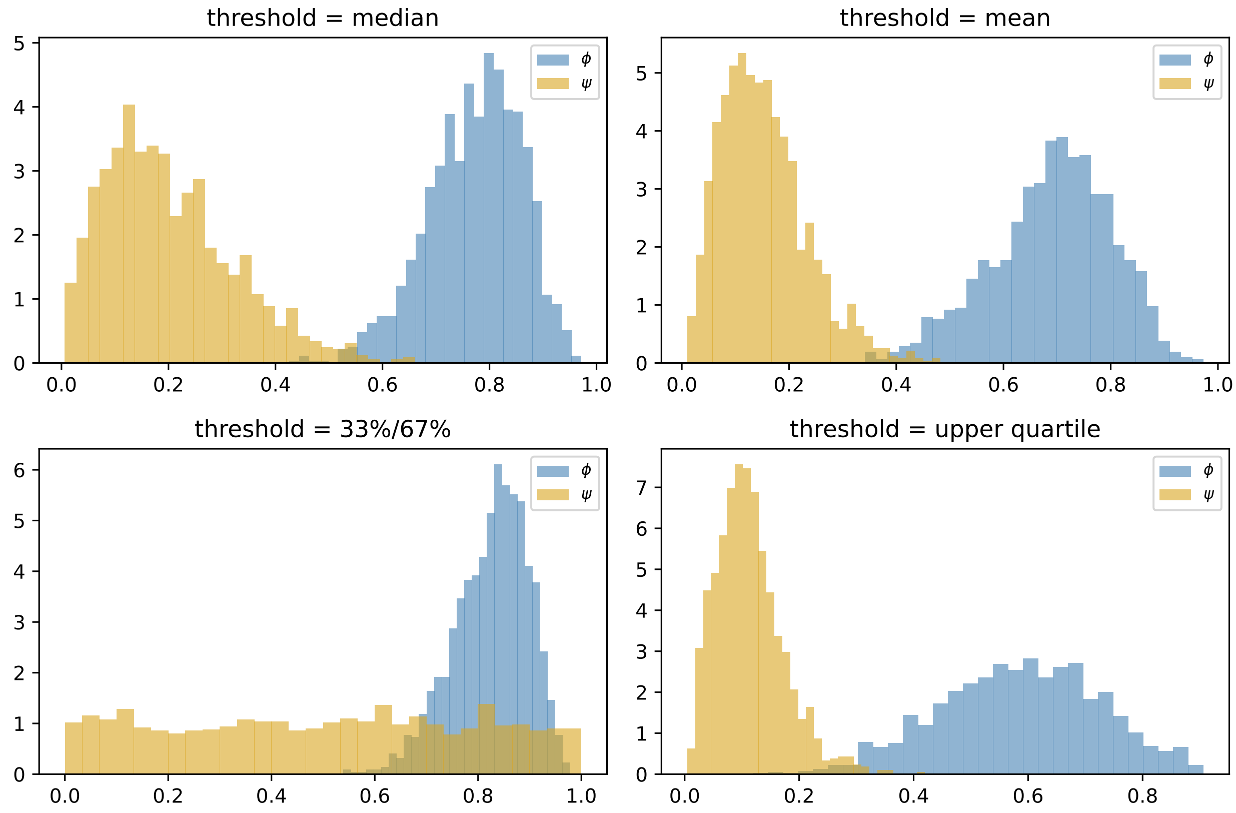

Figure S7: Posterior of $\phi$ (blue) and $\psi$ (gold) under four binarization thresholds. The persistence is robust to threshold choice except in the upper-quartile case where the binary signal becomes sparse.

Listing 1: Neighborhood, $\theta$-grid, and forward simulation

```
# ==============================================================
# sabm_core.R
# Core utilities for the statistical agent-based model (SABM).
#
# Implements the hierarchical Bayesian binary spatio-temporal model:
#
#   y_{i,t} | theta_{i,t} ~ Bern(theta_{i,t})
#
#   theta_{i,t} = y_{i,t-1} * phi
#               + (1 - y_{i,t-1}) * I_{N,i,t-1} * pbar_{i,t}
#               + (1 - I_{N,i,t-1}) * psi
#
# where pbar_{i,t} = 1 - exp( yN_{i,t-1}' log(1 - p) ).
#
# This file provides:
#   - neighborhood index construction (Moore 8-neighborhood with
#     direction-indexed offsets)
#   - the function `theta_grid()` returning theta for every cell
#   - the function `simulate_abm()` for one-step-ahead simulation
#
# Conventions
# -----------
```

```r
# Grid is nrow x ncol. Cells are addressed (r, c) with r = row index
# (south-to-north if you like) and c = column index (west-to-east).
# Directions are numbered 1..8 in the order:
#      1 = N  (r+1, c  )
#      2 = NE (r+1, c+1)
#      3 = E  (r  , c+1)
#      4 = SE (r-1, c+1)
#      5 = S  (r-1, c  )
#      6 = SW (r-1, c-1)
#      7 = W  (r  , c-1)
#      8 = NW (r+1, c-1)
# Offsets are unit cell-widths. The Euclidean distances used in the
# gradient formula are 1 for cardinal and sqrt(2) for diagonal.
# ===============================================================

#' Direction offsets and distances for an 8-neighborhood (Moore).
#'
#' @return list with components:
#'   dr, dc : integer vectors of row/column offsets (length 8)
#'   dist   : Euclidean distance to each neighbor (length 8)
#'   unit   : 2 x 8 matrix of unit vectors in (col, row) = (x, y)
direction_table <- function() {
  dr <- c( 1,  1,  0, -1, -1, -1,  0,  1)   # N NE E SE S SW W NW
  dc <- c( 0,  1,  1,  1,  0, -1, -1, -1)
  d  <- sqrt(dr^2 + dc^2)
  unit <- rbind(dc / d, dr / d)             # (x, y)
  list(dr = dr, dc = dc, dist = d, unit = unit)
}

#' Build a list whose i-th element is the linear indices of the
#' 8-neighbors of cell i, ordered by direction (NA where off-grid).
#'
#' @param nrow,ncol grid dimensions
build_neighborhood <- function(nrow, ncol) {
  dt <- direction_table()
  N  <- nrow * ncol
  nbr <- matrix(NA_integer_, nrow = N, ncol = 8)
  for (r in seq_len(nrow)) {
    for (c in seq_len(ncol)) {
      i <- (c - 1) * nrow + r
      for (j in seq_len(8)) {
        rr <- r + dt$dr[j]
        cc <- c + dt$dc[j]
        if (rr >= 1 && rr <= nrow && cc >= 1 && cc <= ncol) {
          nbr[i, j] <- (cc - 1) * nrow + rr
        }
      }
    }
  }
  list(nbr = nbr, dt = dt, dim = c(nrow, ncol))
}

#' For a binary occupancy vector y_prev, compute the 0/1 occupancy of
#' each direction-j neighbor of each cell. Cells with no neighbor in
#' direction j (i.e. off-grid) are treated as unoccupied.
#'
#' @return N x 8 matrix yN, where yN[i, j] = y_prev[nbr[i, j]] or 0.
neighbor_occupancy <- function(y_prev, nbr) {
  N <- length(y_prev)
```

```r
  yN <- matrix(0, N, 8)
  for (j in 1:8) {
    idx <- nbr[, j]
    ok  <- !is.na(idx)
    yN[ok, j] <- y_prev[idx[ok]]
  }
  yN
}

#' Compute theta for every cell given the previous occupancy y_prev,
#' persistence phi, long-distance dispersal psi, and dispersal
#' probabilities p.
#'
#' @param p either a length-8 vector (stationary) or an N x 8 matrix
#'   (nonstationary, p_i for each cell).
theta_grid <- function(y_prev, phi, psi, p, nbr) {
  yN <- neighbor_occupancy(y_prev, nbr)        # N x 8
  IN <- as.integer(rowSums(yN) > 0)            # at least one neighbor

  # pbar_{i,t} = 1 - exp( yN_i' log(1 - p) )
  if (is.matrix(p)) {
    log1mp <- log(pmax(1 - p, .Machine$double.eps))   # N x 8
    pbar   <- 1 - exp(rowSums(yN * log1mp))
  } else {
    log1mp <- log(pmax(1 - p, .Machine$double.eps))   # length 8
    pbar   <- 1 - exp(yN %*% log1mp)
    pbar   <- as.numeric(pbar)
  }

  # The three regimes are disjoint, so distribute (1 - y_prev) across
  # the IN-conditional terms. (The published formula sometimes drops
  # this factor; we restore it to keep theta in [0, 1].)
  y_prev * phi +
    (1 - y_prev) *  IN       * pbar +
    (1 - y_prev) * (1 - IN) * psi
}

#' One step forward simulation: draw y_t given y_{t-1} and parameters.
step_abm <- function(y_prev, phi, psi, p, nbr) {
  theta <- theta_grid(y_prev, phi, psi, p, nbr)
  rbinom(length(theta), 1, pmin(pmax(theta, 0), 1))
}

#' Full forward simulation over T time steps starting from y0.
#'
#' @param y0 length-N binary vector of initial occupancy.
#' @return T x N integer matrix Y with Y[1, ] = y0.
simulate_abm <- function(y0, Tn, phi, psi, p, nbr) {
  N <- length(y0)
  Y <- matrix(0L, Tn, N)
  Y[1, ] <- y0
  for (t in 2:Tn) {
    Y[t, ] <- step_abm(Y[t - 1, ], phi, psi, p, nbr)
  }
  Y
}
```

#### S4.3 R source: R/sabm_mcmc.R

Listing 2: MCMC for the stationary and nonstationary SABM

```r
# ================================================================
# sabm_mcmc.R
# MCMC inference for the SABM.
#
# Full-conditionals used:
#
#   phi | . ~ Beta( sum_t sum_{i in A_phi,t-1} y_{i,t} + alpha_phi,
#                   sum_t sum_{i in A_phi,t-1} (1 - y_{i,t}) + beta_phi )
#
#   psi | . ~ Beta( sum_t sum_{i in A_psi,t-1} y_{i,t} + alpha_psi,
#                   sum_t sum_{i in A_psi,t-1} (1 - y_{i,t}) + beta_psi )
#
# where
#   A_phi,t-1  = { i : y_{i,t-1} = 1 }
#   A_psi,t-1  = { i : y_{i,t-1} = 0  AND  I_{N,i,t-1} = 0 }.
#
# For the stationary direction-vector p (length 8) we use log(a) ~ N(mu_a,
   Sigma_a)
# with p ~ Dir(a). The Dirichlet prior on p is conjugate only if the
# neighborhood likelihood factors into per-direction Bernoullis, which it
# does NOT (pbar is 1 - prod(1 - p_j)^{yN_j}). We therefore use Metropolis-
# Hastings on log(a) with a random-walk normal proposal.
#
# For the nonstationary model we sample beta with a Gaussian random walk
# (suitability hyperparameters sigma_alpha, theta_alpha are held fixed
# in this reference implementation; extending to slice / MH updates is
# straightforward).
# ================================================================

if (!exists("theta_grid", mode = "function")) source("R/sabm_core.R")
if (!exists("build_p_nonstationary", mode = "function"))
  source("R/sabm_nonstationary.R")

# ---------- helpers ---------------------------------------------

#' Sum y_{i,t} and (1 - y_{i,t}) over the "persistence" cells
#' (cells occupied at the previous time step).
sufficient_phi <- function(Y) {
  Tn <- nrow(Y)
  s1 <- 0; s0 <- 0
  for (t in 2:Tn) {
    mask <- Y[t - 1, ] == 1
    s1 <- s1 + sum(Y[t, mask] == 1)
    s0 <- s0 + sum(Y[t, mask] == 0)
  }
  c(s1, s0)
}

#' Sum over the "long-distance" cells (unoccupied AND no occupied neighbor).
sufficient_psi <- function(Y, nbr) {
  Tn <- nrow(Y); s1 <- 0; s0 <- 0
  for (t in 2:Tn) {
    yN <- neighbor_occupancy(Y[t - 1, ], nbr)
    IN <- rowSums(yN) > 0
    mask <- (Y[t - 1, ] == 0) & (!IN)
```

```r
    s1 <- s1 + sum(Y[t, mask] == 1)
    s0 <- s0 + sum(Y[t, mask] == 0)
  }
  c(s1, s0)
}

#' Log-likelihood of the full dataset given parameters; useful for the
#' p / beta MH steps.
loglik_abm <- function(Y, phi, psi, p, nbr) {
  Tn <- nrow(Y); ll <- 0
  for (t in 2:Tn) {
    th <- theta_grid(Y[t - 1, ], phi, psi, p, nbr)
    th <- pmin(pmax(th, 1e-12), 1 - 1e-12)
    ll <- ll + sum(Y[t, ] * log(th) + (1 - Y[t, ]) * log(1 - th))
  }
  ll
}

# ---------- stationary MCMC -------------------------------------

#' MCMC for the stationary anisotropic SABM.
#'
#' @param Y          T x N binary matrix of observations.
#' @param nbr        output of build_neighborhood().
#' @param n_iter     total MCMC iterations.
#' @param prior      list with alpha_phi, beta_phi, alpha_psi, beta_psi,
#'                   mu_a (length 8), Sigma_a (8x8) for log(a).
#' @param init       list with phi, psi, log_a.
#' @param prop_sd    RW step sd for log(a).
mcmc_stationary <- function(Y, nbr, n_iter = 2000,
                            prior = NULL, init = NULL,
                            prop_sd = 0.1, verbose = TRUE) {

  if (is.null(prior)) prior <- list(
    alpha_phi = 1, beta_phi = 1,
    alpha_psi = 1, beta_psi = 1,
    mu_a = rep(0, 8), Sigma_a = diag(4, 8))
  if (is.null(init)) init <- list(
    phi = 0.5, psi = 0.01, log_a = rep(0, 8))

  inv_Sigma_a <- solve(prior$Sigma_a)

  s_phi <- sufficient_phi(Y)
  s_psi <- sufficient_psi(Y, nbr)

  phi    <- init$phi
  psi    <- init$psi
  log_a  <- init$log_a
  a      <- exp(log_a)
  p      <- a / sum(a)                    # Dirichlet mean
  ll     <- loglik_abm(Y, phi, psi, p, nbr)

  out_phi <- numeric(n_iter)
  out_psi <- numeric(n_iter)
  out_p   <- matrix(0, n_iter, 8)
  out_ll  <- numeric(n_iter)
  accept  <- 0

  for (it in seq_len(n_iter)) {
```

```r

    # ---- phi: exact Beta full-conditional ----------------------
    phi <- rbeta(1, s_phi[1] + prior$alpha_phi,
                    s_phi[2] + prior$beta_phi)

    # ---- psi: exact Beta full-conditional ----------------------
    psi <- rbeta(1, s_psi[1] + prior$alpha_psi,
                    s_psi[2] + prior$beta_psi)

    # ---- log(a): Gaussian RW MH on the marginal likelihood -----
    log_a_star <- log_a + rnorm(8, 0, prop_sd)
    a_star <- exp(log_a_star)
    p_star <- a_star / sum(a_star)
    ll_star <- loglik_abm(Y, phi, psi, p_star, nbr)

    d  <- log_a    - prior$mu_a
    ds <- log_a_star - prior$mu_a
    lp     <- -0.5 * as.numeric(t(d)  %*% inv_Sigma_a %*% d)
    lp_str <- -0.5 * as.numeric(t(ds) %*% inv_Sigma_a %*% ds)

    log_alpha <- (ll_star + lp_str) - (ll + lp)
    if (log(runif(1)) < log_alpha) {
      log_a <- log_a_star; a <- a_star; p <- p_star; ll <- ll_star
      accept <- accept + 1
    } else {
      # recompute ll with updated phi/psi (they changed above)
      ll <- loglik_abm(Y, phi, psi, p, nbr)
    }

    out_phi[it] <- phi
    out_psi[it] <- psi
    out_p[it, ] <- p
    out_ll[it]  <- ll

    if (verbose && it %% max(1, n_iter %/% 10) == 0)
      cat(sprintf("  iter %5d  phi=%.3f psi=%.4f  acc(p)=%.2f  ll=%.1f\n",
                  it, phi, psi, accept / it, ll))
  }

  list(phi = out_phi, psi = out_psi, p = out_p, loglik = out_ll,
       accept_p = accept / n_iter)
}

# ---------- nonstationary MCMC ----------------------------------

#' MCMC for the nonstationary SABM. beta is updated with a Gaussian
#' RW MH proposal; phi, psi keep their exact Beta full-conditionals.
#' sigma_alpha, theta_alpha, c_scale are treated as fixed hyperparams
#' (extend with their own MH steps as needed).
mcmc_nonstationary <- function(Y, X, nbr, Dmat,
                               sigma_alpha = 1, theta_alpha = 5,
                               c_scale = 1,
                               n_iter = 2000, prior = NULL,
                               init = NULL, prop_sd_beta = 0.05,
                               verbose = TRUE) {

  P <- ncol(X)
  if (is.null(prior)) prior <- list(
    alpha_phi = 1, beta_phi = 1,
```

```r
    alpha_psi = 1, beta_psi = 1,
    mu_beta = rep(0, P), Sigma_beta = diag(10, P))
  if (is.null(init)) init <- list(
    phi = 0.5, psi = 0.01, beta = rep(0, P))

  inv_Sigma_beta <- solve(prior$Sigma_beta)

  s_phi <- sufficient_phi(Y)
  s_psi <- sufficient_psi(Y, nbr)

  phi  <- init$phi; psi <- init$psi; beta <- init$beta
  bp   <- build_p_nonstationary(X, beta, sigma_alpha, theta_alpha,
                                Dmat, nbr, c_scale)
  Pmat <- bp$P
  ll   <- loglik_abm(Y, phi, psi, Pmat, nbr)

  out_phi  <- numeric(n_iter)
  out_psi  <- numeric(n_iter)
  out_beta <- matrix(0, n_iter, P)
  out_ll   <- numeric(n_iter)
  accept   <- 0

  for (it in seq_len(n_iter)) {
    phi <- rbeta(1, s_phi[1] + prior$alpha_phi,
                    s_phi[2] + prior$beta_phi)
    psi <- rbeta(1, s_psi[1] + prior$alpha_psi,
                    s_psi[2] + prior$beta_psi)

    beta_star <- beta + rnorm(P, 0, prop_sd_beta)
    bp_star   <- build_p_nonstationary(X, beta_star, sigma_alpha,
                                       theta_alpha, Dmat, nbr,
                                       c_scale, alpha = bp$alpha)
    # alpha is held fixed here to integrate out the spatial residual
    # implicitly through the fixed realization; for a fully Bayesian
    # treatment of alpha, add a Gaussian-process update step.
    ll_star <- loglik_abm(Y, phi, psi, bp_star$P, nbr)

    d  <- beta      - prior$mu_beta
    ds <- beta_star - prior$mu_beta
    lp     <- -0.5 * as.numeric(t(d)  %*% inv_Sigma_beta %*% d)
    lp_str <- -0.5 * as.numeric(t(ds) %*% inv_Sigma_beta %*% ds)

    log_alpha <- (ll_star + lp_str) - (ll + lp)
    if (log(runif(1)) < log_alpha) {
      beta <- beta_star; bp <- bp_star; Pmat <- bp$P; ll <- ll_star
      accept <- accept + 1
    } else {
      ll <- loglik_abm(Y, phi, psi, Pmat, nbr)
    }

    out_phi[it]   <- phi
    out_psi[it]   <- psi
    out_beta[it,] <- beta
    out_ll[it]    <- ll

    if (verbose && it %% max(1, n_iter %/% 10) == 0)
      cat(sprintf("  iter %5d  phi=%.3f psi=%.4f acc(beta)=%.2f ll=%.1f\n",
                  it, phi, psi, accept / it, ll))
  }
```

```

  list(phi = out_phi, psi = out_psi, beta = out_beta, loglik = out_ll,
       accept_beta = accept / n_iter, alpha = bp$alpha)
}
```

#### S4.4 R source: R/sabm_nonstationary.R

Listing 3: Nonstationary kernel from suitability gradients

```
# ==============================================================
# sabm_nonstationary.R
# Anisotropic nonstationary SABM: dispersal probabilities depend on
# habitat-suitability gradients.
#
# Suitability surface:
#   alpha = X beta + epsilon,    epsilon ~ N(0, Sigma_alpha)
#   Sigma_alpha = sigma_alpha^2 * exp( -D / theta_alpha )
#
# Directional gradient at cell i in direction j:
#   a_tilde_{i,j} = ( alpha_{N_{j,i}} - alpha_i ) / d_{N_{j,i}}
#
# Concentration vector and transition probabilities:
#   a_i = c * Phi( a_tilde_i )            (componentwise, Phi = N(0,1) CDF)
#   p_i = a_i / sum(a_i)                  (mean of Dir(a_i))
#
# In sampling mode one can instead draw p_i ~ Dir(a_i).
# ==============================================================

source_if_needed <- function(f) {
  if (!exists("direction_table", mode = "function")) source(f)
}
source_if_needed("R/sabm_core.R")

#' Pairwise Euclidean distance matrix on the grid.
grid_distance_matrix <- function(nrow, ncol) {
  rs <- rep(seq_len(nrow), times = ncol)
  cs <- rep(seq_len(ncol), each  = nrow)
  coords <- cbind(cs, rs)               # (x, y)
  as.matrix(dist(coords))
}

#' Draw a suitability surface alpha = X %*% beta + epsilon with an
#' exponential covariance epsilon ~ N(0, sigma^2 exp(-D / theta)).
draw_suitability <- function(X, beta, sigma_alpha, theta_alpha, Dmat) {
  mu     <- as.numeric(X %*% beta)
  Sigma  <- sigma_alpha^2 * exp(-Dmat / theta_alpha)
  # jitter for numerical PD
  L <- tryCatch(chol(Sigma + diag(1e-8, nrow(Sigma))),
                error = function(e) chol(Sigma + diag(1e-4, nrow(Sigma))))
  z <- rnorm(length(mu))
  as.numeric(mu + crossprod(L, z))
}

#' Compute directional gradients a_tilde_{i,j} for every cell and
#' direction. Off-grid neighbors are filled with 0 (no gradient).
directional_gradients <- function(alpha, nbr) {
  N <- length(alpha)
```

```r
  dt <- direction_table()
  a_tilde <- matrix(0, N, 8)
  for (j in 1:8) {
    idx <- nbr[, j]
    ok  <- !is.na(idx)
    a_tilde[ok, j] <- (alpha[idx[ok]] - alpha[ok]) / dt$dist[j]
  }
  a_tilde
}

#' Map gradients to concentration vectors via a_i = c * Phi(a_tilde_i).
#' Off-grid directions (originally 0) get c * Phi(0) = c/2, which is a
#' sensible neutral baseline; if you prefer to zero them out, set them
#' to NA in ‘nbr‘ upstream and they will be masked.
concentration_from_gradients <- function(a_tilde, c_scale = 1.0) {
  c_scale * pnorm(a_tilde)
}

#' Convert concentration matrix a (N x 8) to mean transition probs p_i.
mean_transition <- function(a) {
  s <- rowSums(a)
  s[s == 0] <- 1
  a / s
}

#' Draw p_i ~ Dir(a_i) row-by-row (used if you want stochastic p in
#' simulation rather than its Dirichlet mean).
sample_transition <- function(a) {
  N <- nrow(a)
  P <- matrix(0, N, 8)
  for (i in seq_len(N)) {
    ai <- pmax(a[i, ], 1e-6)
    g  <- rgamma(8, shape = ai, rate = 1)
    P[i, ] <- g / sum(g)
  }
  P
}

#' Convenience: full pipeline from beta / sigma / theta to p matrix.
build_p_nonstationary <- function(X, beta, sigma_alpha, theta_alpha,
                                  Dmat, nbr, c_scale = 1.0,
                                  alpha = NULL, sample = FALSE) {
  if (is.null(alpha))
    alpha <- draw_suitability(X, beta, sigma_alpha, theta_alpha, Dmat)
  a_tilde <- directional_gradients(alpha, nbr)
  a       <- concentration_from_gradients(a_tilde, c_scale)
  P       <- if (sample) sample_transition(a) else mean_transition(a)
  list(alpha = alpha, a_tilde = a_tilde, a = a, P = P)
}
```

#### S4.5 R source: R/sabm_pde.R

Listing 4: Advection–diffusion PDE solver and Lagrangian sanity check

```r
# ==============================================================
# sabm_pde.R
# PDE scaling: the Lagrangian recurrence for occupancy probability
```

```r
# converges (in the small-step limit) to a 2D advection-diffusion
# equation
#
#   d pbar / d t = -delta_1 d pbar/ds1 - delta_2 d pbar/ds2
#                  + D_1 d^2 pbar/ds1^2 + D_2 d^2 pbar/ds2^2
#
# Drift and dispersion are the first- and second-direction moments
# of the dispersal kernel p (averaged across the 8 cardinal/diagonal
# unit vectors):
#
#   delta_k = sum_j p_j * u_{j,k}
#   D_k     = (1/2) * sum_j p_j * u_{j,k}^2
#
# where u_j is the unit displacement of direction j (cell widths per
# time step). For a strongly clustered front pbar is approximately the
# average occupancy probability.
# ==============================================================

if (!exists("direction_table", mode = "function")) source("R/sabm_core.R")

#' Drift / dispersion from a (length-8) probability vector p.
pde_coefficients <- function(p) {
  dt <- direction_table()
  u  <- dt$unit                              # 2 x 8 (x, y)
  delta <- u %*% p                           # 2 x 1
  D     <- 0.5 * (u^2 %*% p)                 # 2 x 1
  list(delta = as.numeric(delta), D = as.numeric(D))
}

#' Stable explicit finite-difference time-step length for an
#' advection-diffusion solver on a unit grid.
pde_dt_max <- function(delta, D, h = 1, safety = 0.4) {
  # CFL for advection: |delta| dt / h <= 1
  # Stability for diffusion: 2 D dt / h^2 <= 1
  dt_adv <- if (max(abs(delta)) > 0) h / max(abs(delta)) else Inf
  dt_dif <- if (max(D) > 0) h^2 / (2 * max(D)) else Inf
  safety * min(dt_adv, dt_dif, 1)
}

#' Solve the advection-diffusion PDE forward in time on a grid with
#' periodic boundaries. Times are in the same units as the ABM steps
#' (one ABM step = one PDE time unit), so the solver subdivides each
#' unit step into n_sub stable explicit substeps.
#'
#' @param u0 nrow x ncol matrix of initial occupancy probability
#' @param p  length-8 dispersal probability vector
#' @param Tn number of integer time steps to advance
#' @return list with $u (Tn x nrow x ncol array) and $coef
solve_pde <- function(u0, p, Tn) {
  nrow <- nrow(u0); ncol <- ncol(u0)
  coef <- pde_coefficients(p)
  dt_max <- pde_dt_max(coef$delta, coef$D)
  n_sub <- max(1, ceiling(1 / dt_max))
  dt    <- 1 / n_sub

  delta_x <- coef$delta[1]; delta_y <- coef$delta[2]
  Dx <- coef$D[1];          Dy <- coef$D[2]

  # shift(M, drow, dcol) returns a matrix whose entry (r, c) equals
```

```r
  # M[r - drow, c - dcol] (mod size). So shift(M, 0, 1)[r, c] = M[r, c - 1]
  # is M shifted one column to the right (value at c comes from the left).
  shift <- function(M, drow, dcol) {
    nr <- nrow(M); nc <- ncol(M)
    r_idx <- ((seq_len(nr) - 1 - drow) %% nr) + 1
    c_idx <- ((seq_len(nc) - 1 - dcol) %% nc) + 1
    M[r_idx, c_idx, drop = FALSE]
  }

  U <- array(0, dim = c(Tn, nrow, ncol))
  U[1, , ] <- u0
  u <- u0
  for (t in 2:Tn) {
    for (s in seq_len(n_sub)) {
      # Central differences on a unit grid. du/dx at (r, c) =
      # (u(r, c+1) - u(r, c-1)) / 2 = (shift(u, 0, -1) - shift(u, 0, 1)) / 2.
      dudx <- (shift(u, 0, -1) - shift(u, 0,  1)) / 2
      dudy <- (shift(u, -1, 0) - shift(u, 1,  0)) / 2
      d2x  <-  shift(u, 0,  1) - 2 * u + shift(u, 0, -1)
      d2y  <-  shift(u, 1,  0) - 2 * u + shift(u, -1, 0)
      rhs  <- -delta_x * dudx - delta_y * dudy + Dx * d2x + Dy * d2y
      u <- u + dt * rhs
      u <- pmin(pmax(u, 0), 1)
    }
    U[t, , ] <- u
  }
  list(u = U, coef = coef, n_sub = n_sub)
}

#' Run many independent ABM trajectories and return the cell-wise mean
#' occupancy, which approximates pbar(s, t) when used as a sanity check
#' on the full binary-occupancy dynamics.
abm_mean_field <- function(y0, Tn, phi, psi, p, nbr, n_rep = 200) {
  N <- length(y0)
  Y_mean <- matrix(0, Tn, N)
  for (r in seq_len(n_rep)) {
    Yr <- simulate_abm(y0, Tn, phi, psi, p, nbr)
    Y_mean <- Y_mean + Yr
  }
  Y_mean / n_rep
}

#' Simulate the position distribution of a SINGLE Lagrangian agent on
#' the grid. At each step the agent moves to neighbor j with prob p_j
#' (a length-8 vector; sum(p) is assumed 1). Off-grid moves are
#' rejected (the agent stays put), so the boundary acts reflecting.
#' Averaging over many trajectories yields the discrete approximation
#' to the advection-diffusion PDE's solution pbar(s, t).
simulate_agent_field <- function(start_rc, nrow, ncol, Tn, p,
                                 n_rep = 1000) {
  dt <- direction_table()
  pn <- p / sum(p)
  field <- array(0, dim = c(Tn, nrow, ncol))
  for (r in seq_len(n_rep)) {
    rr <- start_rc[1]; cc <- start_rc[2]
    field[1, rr, cc] <- field[1, rr, cc] + 1
    for (t in 2:Tn) {
      j <- sample.int(8, 1, prob = pn)
      nr <- rr + dt$dr[j]; nc <- cc + dt$dc[j]
```

```r
      if (nr >= 1 && nr <= nrow && nc >= 1 && nc <= ncol) {
        rr <- nr; cc <- nc
      }
      field[t, rr, cc] <- field[t, rr, cc] + 1
    }
  }
  field / n_rep
}

#' Compare the agent position distribution from Lagrangian simulation
#' to the advection-diffusion PDE solution. Both share the same drift
#' and dispersion coefficients derived from p.
compare_agent_pde <- function(start_rc, nrow, ncol, Tn, p,
                              n_rep = 2000) {
  u0 <- matrix(0, nrow, ncol)
  u0[start_rc[1], start_rc[2]] <- 1
  pde <- solve_pde(u0, p, Tn)
  agt <- simulate_agent_field(start_rc, nrow, ncol, Tn, p, n_rep)
  rmse <- sqrt(mean((agt - pde$u)^2))
  list(agent = agt, pde = pde$u, rmse = rmse, coef = pde$coef)
}

#' Compare the binary ABM mean-occupancy field to the advection-
#' diffusion PDE. The agreement is best when p is small (the PDE is a
#' small-step linearization of pbar = 1 - exp(-yN' p)).
compare_abm_pde <- function(y0_mat, Tn, phi, psi, p, nbr, n_rep = 200) {
  N <- length(y0_mat)
  y0 <- as.integer(as.vector(y0_mat))
  abm <- abm_mean_field(y0, Tn, phi, psi, p, nbr, n_rep = n_rep)
  pde <- solve_pde(y0_mat, p, Tn)
  nrow <- nrow(y0_mat); ncol <- ncol(y0_mat)
  abm_arr <- array(abm, dim = c(Tn, nrow, ncol))
  rmse <- sqrt(mean((abm_arr - pde$u)^2))
  list(abm = abm_arr, pde = pde$u, rmse = rmse, coef = pde$coef)
}
```

##### S4.6 R source: R/sabm_temporal.R

Listing 5: Temporal-only special case ($N = 1$) used for univariate series

```r
# ==============================================================
# sabm_temporal.R
# Temporal-only special case of the SABM, used for univariate
# binary time series. With N = 1 there is no neighborhood, so
# the indicator I_{N,i,t-1} is identically 0 and the data model
# collapses to a 2-state Markov chain:
#
#   theta_t = y_{t-1} * phi + (1 - y_{t-1}) * psi.
#
# Both phi and psi retain conjugate Beta full-conditionals.
# This file provides the conjugate Gibbs sampler used for the
# lynx and hare data application.
# ==============================================================

#' Sufficient statistics for the temporal SABM. Multiple disjoint
#' time-series segments (e.g. before/after a gap) can be combined
#' by passing a list of binary vectors `y_list`.
```

```
sufficient_temporal <- function(y_list) {
  if (!is.list(y_list)) y_list <- list(y_list)
  s_phi <- c(0, 0); s_psi <- c(0, 0)
  for (y in y_list) {
    if (length(y) < 2) next
    for (t in 2:length(y)) {
      if (y[t - 1] == 1) {
        if (y[t] == 1) s_phi[1] <- s_phi[1] + 1
        else           s_phi[2] <- s_phi[2] + 1
      } else {
        if (y[t] == 1) s_psi[1] <- s_psi[1] + 1
        else           s_psi[2] <- s_psi[2] + 1
      }
    }
  }
  list(phi = s_phi, psi = s_psi)
}

#' Conjugate Gibbs sampler. Returns posterior draws for phi, psi.
gibbs_temporal <- function(y_list, n_iter = 5000,
                           prior = list(a_phi = 1, b_phi = 1,
                                        a_psi = 1, b_psi = 1)) {
  s <- sufficient_temporal(y_list)
  phi <- rbeta(n_iter, s$phi[1] + prior$a_phi,
                       s$phi[2] + prior$b_phi)
  psi <- rbeta(n_iter, s$psi[1] + prior$a_psi,
                       s$psi[2] + prior$b_psi)
  list(phi = phi, psi = psi, suff = s)
}

#' Posterior predictive: simulate binary sequences from the
#' draws and report cell-wise occupancy frequency.
posterior_predictive <- function(fit, T_pred, y0 = 0L,
                                 n_rep = NULL) {
  if (is.null(n_rep)) n_rep <- length(fit$phi)
  ix <- sample(seq_along(fit$phi), n_rep, replace = TRUE)
  Y  <- matrix(0L, n_rep, T_pred)
  Y[, 1] <- y0
  for (r in seq_len(n_rep)) {
    phi <- fit$phi[ix[r]]; psi <- fit$psi[ix[r]]
    for (t in 2:T_pred) {
      theta <- if (Y[r, t - 1] == 1) phi else psi
      Y[r, t] <- rbinom(1, 1, theta)
    }
  }
  list(Y = Y, mean = colMeans(Y),
       boom_rate = mean(Y),
       cycle_len = mean(apply(Y, 1, runs_to_cycle)))
}

#' Estimate cycle length from a binary sequence via the average
#' run-length 0->1 transitions.
runs_to_cycle <- function(y) {
  starts <- which(diff(c(0L, y)) == 1L)
  if (length(starts) < 2) return(NA_real_)
  mean(diff(starts))
}

#' Convenience: binarize numeric counts via threshold (default
```

```r
#' median). NA values stay NA.
binarize <- function(x, thresh = NULL) {
  if (is.null(thresh)) thresh <- median(x, na.rm = TRUE)
  ifelse(is.na(x), NA_integer_, as.integer(x > thresh))
}

#' Break a possibly-NA binary series into contiguous non-NA runs.
contiguous_segments <- function(y) {
  segs <- list(); cur <- integer(0)
  for (v in y) {
    if (is.na(v)) {
      if (length(cur) >= 2) segs[[length(segs) + 1]] <- cur
      cur <- integer(0)
    } else cur <- c(cur, v)
  }
  if (length(cur) >= 2) segs[[length(segs) + 1]] <- cur
  segs
}
```

#### S4.7 Application driver: validate_lynx_hare.R

Listing 6: Driver for the lynx–hare application

```r
# ==============================================================
# validate_lynx_hare.R
# Real-data validation of the SABM using the Hudson's Bay Company
# lynx-hare pelt records.
#
#   * Single-cell fit on the full lynx series (1821-1934, n = 114)
#     -- tests persistence phi and long-distance psi alone.
#   * Two-cell fit on the joint hare/lynx series (1900-1920, n = 21)
#     -- introduces the directional kernel via a 1x2 spatial grid
#     so that the two species act as each other's only neighbor.
#   * Posterior predictive check + held-out one-step-ahead accuracy.
#
# Run from the repository root:
#     Rscript validate_lynx_hare.R
# ==============================================================

source("R/sabm_core.R")
source("R/sabm_mcmc.R")

set.seed(20260601)

# ---------------- Hudson Bay hare-lynx data ---------------------
# Annual pelt counts (thousands) from the Hudson's Bay Company.
# Source: MacLulich (1937), as compiled in many introductory texts.
hb <- data.frame(
  year = 1900:1920,
  hare = c(30.0, 47.2, 70.2, 77.4, 36.3, 20.6, 18.1, 21.4, 22.0, 25.4,
           27.1, 40.3, 57.0, 76.6, 52.3, 19.5, 11.2,  7.6, 14.6, 16.2,
           24.7),
  lynx = c( 4.0,  6.1,  9.8, 35.2, 59.4, 41.7, 19.0, 13.0,  8.3,  9.1,
            7.4,  8.0, 12.3, 19.5, 45.7, 51.1, 29.7, 15.8,  9.7, 10.1,
            8.6))

binarize <- function(x, thr = median(x)) as.integer(x > thr)
```

```
y_hare <- binarize(hb$hare)
y_lynx <- binarize(hb$lynx)

cat("Hudson Bay 1900-1920 binarized series:\n")
cat("  hare:", paste(y_hare, collapse = ""), "\n")
cat("  lynx:", paste(y_lynx, collapse = ""), "\n\n")

# ---------------- 1-cell SABM (lynx, long series) ----------------
# datasets::lynx is the 114-year (1821-1934) Canadian Lynx series.
ln <- as.numeric(datasets::lynx)
y_long <- binarize(ln)
T_long <- length(y_long)
nbr1   <- build_neighborhood(1, 1)
Y1     <- matrix(y_long, ncol = 1)

cat("Lynx (1821-1934): n =", T_long,
    "  fraction 'high' =", round(mean(y_long), 2), "\n")

# With one cell, the neighborhood term vanishes (I_N = 0 always), so
# phi and psi reduce to a two-state Markov chain. We use the exact
# Beta full-conditionals directly.
sphi  <- sufficient_phi(Y1)
spsi  <- sufficient_psi(Y1, nbr1)
phi_post <- rbeta(5000, sphi[1] + 1, sphi[2] + 1)
psi_post <- rbeta(5000, spsi[1] + 1, spsi[2] + 1)

cat(sprintf("  phi posterior (lynx)  : mean %.3f  95%% CrI [%.3f, %.3f]\n",
            mean(phi_post),
            quantile(phi_post, 0.025), quantile(phi_post, 0.975)))
cat(sprintf("  psi posterior (lynx)  : mean %.3f  95%% CrI [%.3f, %.3f]\n\n",
            mean(psi_post),
            quantile(psi_post, 0.025), quantile(psi_post, 0.975)))

# Stationary occupancy implied by the two-state Markov chain.
pi_lynx <- mean(psi_post / (1 - phi_post + psi_post))
cat(sprintf("  implied stationary P(high) = %.3f   (data: %.3f)\n\n",
            pi_lynx, mean(y_long)))

# ---------------- 1-cell SABM (hare alone) -----------------------
y_hare_long <- y_hare
Yh    <- matrix(y_hare_long, ncol = 1)
sphih <- sufficient_phi(Yh)
spsih <- sufficient_psi(Yh, nbr1)
phi_h <- rbeta(5000, sphih[1] + 1, sphih[2] + 1)
psi_h <- rbeta(5000, spsih[1] + 1, spsih[2] + 1)
cat(sprintf("Hare (1900-1920, n=21): phi mean %.3f, psi mean %.3f\n\n",
            mean(phi_h), mean(psi_h)))

# ---------------- 2-cell SABM (lynx <-> hare) --------------------
# Place lynx in column 1 and hare in column 2 of a 1x2 grid. Each is
# the other's only on-grid neighbor: only directions 3 (E) and 7 (W)
# of the 8-direction kernel see data, and they encode arrivals
# lynx -> hare (p_3) and hare -> lynx (p_7) respectively.
nbr2 <- build_neighborhood(1, 2)
Y2   <- cbind(y_lynx, y_hare)

cat("Running 2-cell MCMC on the joint lynx/hare series...\n")
fit <- mcmc_stationary(Y2, nbr2, n_iter = 4000,
```

```
                         prop_sd = 0.2, verbose = FALSE)

burn <- 1000
post_mean <- function(x) {
  if (is.matrix(x)) colMeans(x[(burn + 1):nrow(x), , drop = FALSE])
  else               mean(x[(burn + 1):length(x)])
}
phi_hat <- post_mean(fit$phi)
psi_hat <- post_mean(fit$psi)
p_hat   <- post_mean(fit$p)

cat(sprintf("  phi  (persistence)        : %.3f\n", phi_hat))
cat(sprintf("  psi  (long-distance)      : %.3f\n", psi_hat))
cat(sprintf("  p[3] (lynx -> hare arrival): %.3f\n", p_hat[3]))
cat(sprintf("  p[7] (hare -> lynx arrival): %.3f\n", p_hat[7]))
cat(sprintf("  MH acceptance for p        : %.2f\n\n", fit$accept_p))

# ---------------- Posterior predictive check ---------------------
# Forward simulate many replicates from the posterior mean and
# compare to the observed series in two summaries: mean occupancy
# and lag-1 transition counts.
n_rep <- 1000
sim_means <- matrix(0, n_rep, 2)
sim_trans <- array(0, dim = c(n_rep, 2, 4))  # for each cell: 00,01,10,11
for (r in seq_len(n_rep)) {
  y <- Y2[1, ]
  out <- matrix(0, nrow(Y2), 2)
  out[1, ] <- y
  for (t in 2:nrow(Y2))
    out[t, ] <- step_abm(out[t - 1, ], phi_hat, psi_hat, p_hat, nbr2)
  sim_means[r, ] <- colMeans(out)
  for (c in 1:2) {
    pair <- 2 * out[-nrow(out), c] + out[-1, c]   # 0..3
    for (k in 0:3) sim_trans[r, c, k + 1] <- sum(pair == k)
  }
}

obs_means <- colMeans(Y2)
obs_trans <- matrix(0, 2, 4)
for (c in 1:2) {
  pair <- 2 * Y2[-nrow(Y2), c] + Y2[-1, c]
  for (k in 0:3) obs_trans[c, k + 1] <- sum(pair == k)
}

cat("Posterior predictive check (mean occupancy):\n")
cat(sprintf("  lynx: obs %.2f  ppd mean %.2f  95%% [%.2f, %.2f]\n",
            obs_means[1], mean(sim_means[, 1]),
            quantile(sim_means[, 1], 0.025),
            quantile(sim_means[, 1], 0.975)))
cat(sprintf("  hare: obs %.2f  ppd mean %.2f  95%% [%.2f, %.2f]\n\n",
            obs_means[2], mean(sim_means[, 2]),
            quantile(sim_means[, 2], 0.025),
            quantile(sim_means[, 2], 0.975)))

cat("Posterior predictive check (lag-1 transition counts, lynx):\n")
cat("  pattern  obs  ppd mean  (95% interval)\n")
labs <- c("00", "01", "10", "11")
for (k in 1:4)
  cat(sprintf("  %s     %3d   %5.1f    [%2d, %2d]\n",
```

```
                 labs[k], obs_trans[1, k], mean(sim_trans[, 1, k]),
                 floor(quantile(sim_trans[, 1, k], 0.025)),
                 ceiling(quantile(sim_trans[, 1, k], 0.975))))
cat("\nPosterior predictive check (lag-1 transition counts, hare):\n")
for (k in 1:4)
  cat(sprintf("  %s     %3d   %5.1f    [%2d, %2d]\n",
                 labs[k], obs_trans[2, k], mean(sim_trans[, 2, k]),
                 floor(quantile(sim_trans[, 2, k], 0.025)),
                 ceiling(quantile(sim_trans[, 2, k], 0.975))))

# ---------------- One-step-ahead held-out accuracy ---------------
# Fit on the first 16 years and forecast the last 5 by computing,
# for each held-out year, the model's predictive probability of
# y_t = 1 given y_{t-1}, and compare to the observed y_t.
T_all <- nrow(Y2); T_fit <- 16
fit_h <- mcmc_stationary(Y2[1:T_fit, ], nbr2, n_iter = 3000,
                         prop_sd = 0.2, verbose = FALSE)
phi_h2 <- mean(fit_h$phi[(burn + 1):3000])
psi_h2 <- mean(fit_h$psi[(burn + 1):3000])
p_h2   <- colMeans(fit_h$p[(burn + 1):3000, ])

cat("\n1-step-ahead held-out probabilities, years 1916-1920\n")
cat("       y_{t-1}    y_t  P(y_t = 1 | y_{t-1})\n")
brier <- 0; correct <- 0; n_pred <- 0
for (t in (T_fit + 1):T_all) {
  theta <- theta_grid(Y2[t - 1, ], phi_h2, psi_h2, p_h2, nbr2)
  for (c in 1:2) {
    spp <- c("lynx", "hare")[c]
    cat(sprintf("  %4d  %s  %d -> %d   %.3f\n", hb$year[t], spp,
                Y2[t - 1, c], Y2[t, c], theta[c]))
    brier <- brier + (Y2[t, c] - theta[c])^2
    correct <- correct + ((theta[c] > 0.5) == (Y2[t, c] == 1))
    n_pred <- n_pred + 1
  }
}
cat(sprintf("\nBrier score = %.3f   accuracy = %d / %d\n",
            brier / n_pred, correct, n_pred))

saveRDS(list(fit = fit, fit_holdout = fit_h,
             phi_post = phi_post, psi_post = psi_post,
             post_pred_means = sim_means,
             post_pred_trans = sim_trans),
        file = "lynx_hare_results.rds")
cat("\nSaved lynx_hare_results.rds\n")
```
